# Genome-scale transcriptional regulatory network models for the mouse and human striatum predict roles for SMAD3 and other transcription factors in Huntington’s disease

**DOI:** 10.1101/087114

**Authors:** Seth A. Ament, Jocelynn R. Pearl, Robert M. Bragg, Peter J. Skene, Sydney R. Coffey, Dani E. Bergey, Christopher L. Plaisier, Vanessa C. Wheeler, Marcy E. MacDonald, Nitin S. Baliga, Jim Rosinski, Leroy E. Hood, Jeffrey B. Carroll, Nathan D. Price

## Abstract

Transcriptional changes occur presymptomatically and throughout Huntington’s Disease (HD), motivating the study of transcriptional regulatory networks (TRNs) in HD. We reconstructed a genome-scale model for the target genes of 718 TFs in the mouse striatum by integrating a model of the genomic binding sites with transcriptome profiling of striatal tissue from HD mouse models. We identified 48 differentially expressed TF-target gene modules associated with age‐ and *Htt* allele-dependent gene expression changes in the mouse striatum, and replicated many of these associations in independent transcriptomic and proteomic datasets. Strikingly, many of these predicted target genes were also differentially expressed in striatal tissue from human disease. We experimentally validated a key model prediction that SMAD3 regulates HD-related gene expression changes using chromatin immunoprecipitation and deep sequencing (ChIP-seq) of mouse striatum. We found *Htt* allele-dependent changes in the genomic occupancy of SMAD3 and confirmed our model’s prediction that many SMAD3 target genes are down-regulated early in HD. Importantly, our study provides a mouse and human striatal-specific TRN and prioritizes a hierarchy of transcription factor drivers in HD.

## Introduction

Massive changes in gene expression accompany many human diseases, yet we still know relatively little about how specific transcription factors (TFs) mediate these changes. Comprehensive characterization of disease-related transcriptional regulatory networks (TRNs) can clarify potential disease mechanisms and prioritize targets for novel therapeutics. A variety of approaches have been developed to reconstruct interactions between TFs and their target genes, including models focused on reconstructing the physical locations of transcription factor binding (Neph *et al*, 2012; Gerstein *et al*, 2012), as well as computational algorithms utilizing gene co-expression to infer regulatory relationships (Marbach *et al*, 2012; Margolin *et al*, 2006; Bonneau *et al*, 2006; Friedman *et al*, 2000; Huynh-Thu *et al*, 2010; Reiss *et al*, 2015). These approaches have yielded insights into the regulation of a range of biological systems, yet accurate, genome-scale models of mammalian TRNs remain elusive.

Several lines of evidence point to a specific role for transcriptional regulatory changes in Huntington’s disease (HD). HD is a fatal neurodegenerative disease caused by dominant inheritance of a polyglutamine (polyQ)-coding expanded trinucleotide (CAG) repeat in the *HTT* gene (MacDonald *et al*, 1993). Widespread transcriptional changes have been detected in post-mortem brain tissue from HD cases vs. controls (Hodges *et al*, 2006), and transcriptional changes are among the earliest detectable phenotypes in HD mouse models (Luthi-Carter *et al*, 2000; Seredenina & Luthi-Carter, 2012). These transcriptional changes are particularly prominent in the striatum, the most profoundly impacted brain region in HD (Tabrizi *et al*, 2013; Vonsattel *et al*, 1985). Replicable gene expression changes in the striatum of HD patients and HD mouse models include down-regulation of genes related to synaptic function in medium spiny neurons accompanied by up-regulation of genes related to neuroinflammation (Seredenina & Luthi-Carter, 2012).

Some of these transcriptional changes may be directly related to the functions of the HTT protein. Both wildtype and mutant HTT (mHTT) protein have been shown to associate with genomic DNA, and mHTT also interacts with histone modifying enzymes and is associated with changes in chromatin states (Thomas *et al*; Benn *et al*; Seong *et al*, 2010). Wildtype HTT protein has been shown to regulate the activity of some TFs (Zuccato *et al*, 2007). Also, high concentrations of nuclear mHTT aggregates sequester TF and co-factor proteins and interfere with genomic target finding, though it is unknown if this occurs at physiological concentrations of mHTT (Wheeler *et al*, 2000; Shirasaki *et al*, 2012; Li *et al*, 2016). Roles for several TFs in HD have been characterized (Huntingtin interacts with REST/NRSF to modulate the transcription of NRSE-controlled neuronal genes, 2003; Dickey *et al*, 2015; Arlotta *et al*, 2008; Tang *et al*, 2012), but we lack a global model for the relationships between HD-related changes in the activity of specific TFs and the downstream pathological processes that they regulate.

The availability of large transcriptomics datasets related to HD is now making it possible to begin comprehensive network analysis of the disease, particularly in mouse models. Langfelder et al. (Langfelder *et al*, 2016) generated RNA-seq from the striatum of 144 knock-in mice heterozygous for HD mutations and 64 wildtype littermate controls, and they used gene co-expression networks to identify modules of co-expressed genes with altered expression in HD. However, their analyses did not attempt to identify any of the TFs responsible for these gene expression changes.

Here, we investigated the roles of core TFs that are predicted to drive the gene expression changes in Huntington’s disease, using a comprehensive network biology approach. We used a machine learning strategy to reconstruct a genome-scale model for TF-target gene interactions in the mouse striatum, combining publicly available DNase-seq with brain transcriptomics data HD mouse models. We identified 48 core TFs whose predicted target genes were overrepresented among differentially expressed genes in at least five of fifteen conditions defined by a mouse’s age and *Htt* allele, and we replicated the predicted core TFs and differential gene expression associations in multiple datasets from HD mouse models and from HD cases and controls. Based on the gene expression signature of SMAD3 and its predicted target genes, we hypothesized that SMAD3 is a core regulator of early gene expression changes in HD. Using chromatin immunoprecipitation and deep sequencing (ChIP-seq), we demonstrate *Htt*-allele-dependent changes in SMAD3 occupancy and down-regulation of SMAD3 target genes in mouse brain tissue. In conclusion, the results from our TRN analysis and ChIP-seq studies of HD reveal new insights into transcription factor drivers of complex gene expression changes in this neurodegenerative disease.

## Results

### A genome-scale transcriptional regulatory network model of the mouse striatum

We reconstructed a model of TF-target gene interactions in the mouse striatum by integrating information about transcription factor binding sites (TFBSs) with evidence from gene co-expression in the mouse striatum (Fig. 1a).

**Figure 1.**
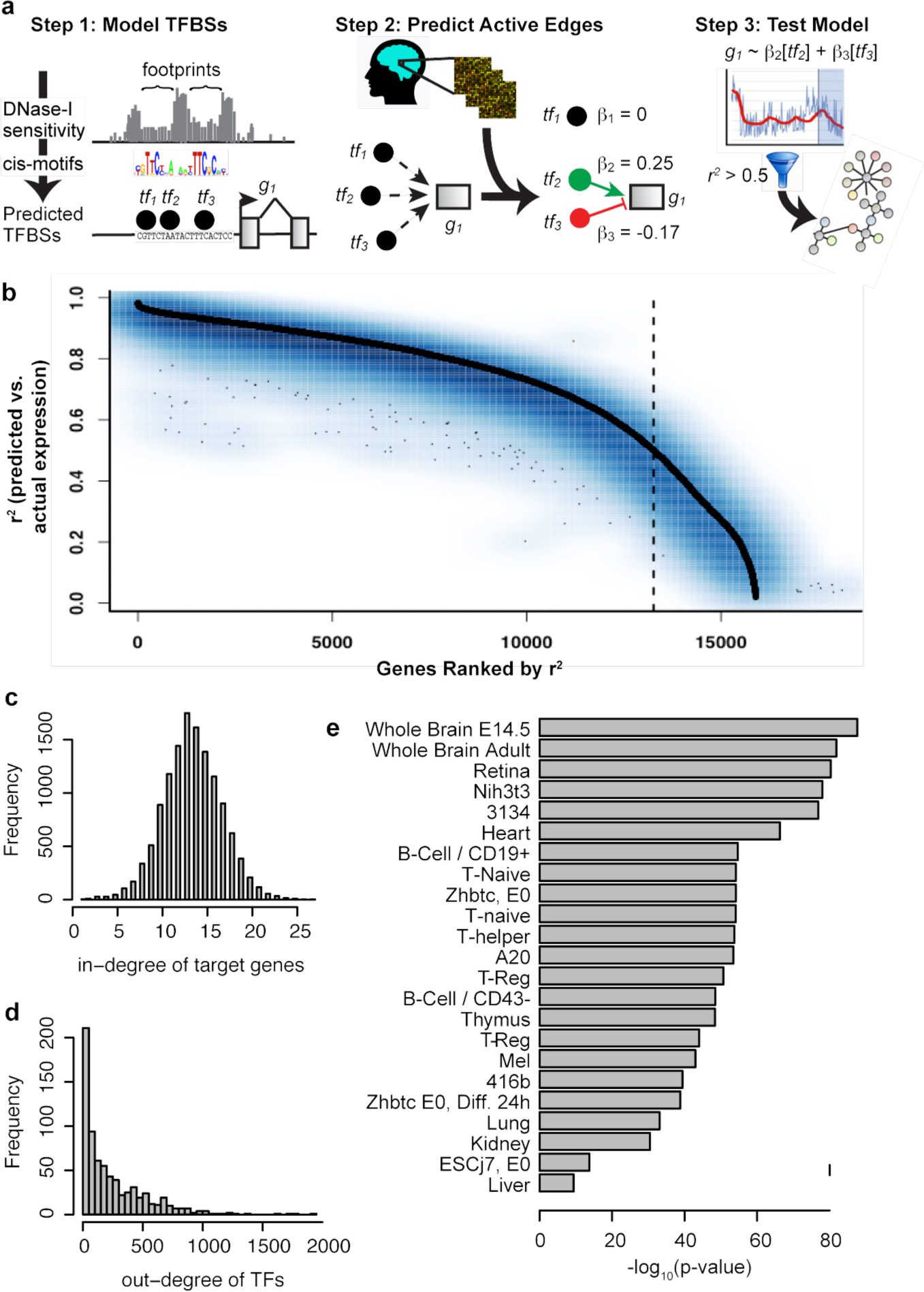
Reconstruction and validation of a transcriptional regulatory network (TRN) model of the mouse striatum. a. Schematic for reconstruction of tissue-specific TRN models by combining information about TF binding sites with evidence from co-expression. b. Training (black) and test set (blue) prediction accuracy for genes in the mouse striatum TRN model. Genes are ordered on the x-axis according to their training set prediction accuracy (r^2^, predicted vs. actual expression). c. Distribution for the number of predicted regulators per target gene. d. Distribution for the number of predicted target genes per TF. e. Enrichments of TF-target gene interactions in the mouse striatum TRN for TFBSs supported by DNase footprints identified in 23 tissues.

We predicted the binding sites for 871 TFs in the mouse genome using digital genomic footprinting. We identified footprints in DNase-seq data from 23 mouse tissues (Yue *et al*, 2014), using Wellington (Piper *et al*, 2013). Footprints are defined as short genomic regions with reduced accessibility to the DNase-I enzyme in at least one tissue. Our goal in combining DNase-seq data from multiple tissues was to reconstruct a single TFBS model that could make useful predictions about TF target genes, even in conditions for which DNase-seq data were not available. We identified 3,242,454 DNase-I footprints. Genomic footprints are often indicative of occupancy by a DNA-binding protein. We scanned these footprints for 2,547 sequence motifs from TRANSFAC (Matys *et al*, 2006), JASPAR (Mathelier *et al*, 2014), UniProbe (Hume *et al*, 2015), and high-throughput SELEX (Jolma *et al*, 2013) to predict binding sites for specific TFs (TFBSs), and we compared these TFBSs to the locations of transcription start sites. We considered a TF to be a potential regulator of a gene if it had at least one binding site within 5kb of that gene’s TSS. We showed previously that a 5kb region upstream and downstream of the TSS maximizes target gene prediction from digital genomic footprinting of the human genome (Plaisier *et al*, 2016).

To assess the accuracy of this TFBS model, we compared our TFBS predictions to ChIP-seq experiments from ENCODE (Yue *et al*, 2014) and ChEA (Lachmann *et al*, 2010) (SI Fig. 2). For 50 of 52 TFs, there was significant overlap between the sets of target genes predicted by our TFBS model vs. ChIP-seq (FDR < 1%). Our TFBS model had a median 78% recall of target genes identified by ChIP-seq, and a median 22% precision. That is, our model identified the majority of true-positive target genes but also made a large number of false-positive predictions. Low precision is expected in this model, since TFs typically occupy only a subset of their binding sites in a given tissue. Nonetheless, low precision indicates a need for additional filtering steps to identify target genes that are relevant in a specific context.

We sought to identify TF-target gene interactions that are active in the mouse striatum, by evaluating gene co-expression patterns in RNA-seq transcriptome profiles from the striatum of 208 mice (Langfelder *et al*, 2016). The general idea is that active regulation of a target gene by a TF is likely to be associated with strong TF-gene co-expression, and TFBSs allow us to identify direct regulatory interactions. This step also removes TFs with low expression: of the 871 TFs with TFBS predictions we retained as potential regulators the 718 TFs that were expressed in the striatum. We fit a regression model to predict the expression of each gene based on the combined expression patterns of TFs with one or more TFBSs ±5kb of that gene’s transcription start site. We used LASSO regularization to select the subset of TFs whose expression patterns together predicted the expression of the target gene. This approach extends several previous regression methods for TRN reconstruction (Friedman *et al*, 2010; Tibshirani, 1996; Bonneau *et al*, 2006; Haury *et al*, 2012) by introducing TFBS-­-based constraints. In preliminary work, we considered a range of LASSO and elastic net (alpha = 0.2, 0.4, 0.6, 0.8, 1.0) regularization penalties and evaluated performance in five-fold cross-validation (see Methods). We selected LASSO based on the highest correlation between prediction accuracy in training vs. test sets.

We validated the predictive accuracy of our TRN model by comparing predicted vs. observed expression levels of each gene. Our model explained >50% of expression variation for 13,009 genes in training data (Fig. 1b). Prediction accuracy in five-fold cross-validation was nearly identical to prediction accuracy in training data. That is, genes whose expression was accurately predicted in the training data were also accurately predicted in the test sets (r=0.94; Fig. 1b). Genes whose expression was not accurately predicted generally had low expression in the striatum (Supplementary Information [SI] Fig. 1). We removed poorly predicted genes, based on their training set accuracy before moving to the test set. The final TRN model contains 13,009 target genes regulated by 718 TFs via 176,518 interactions (SI Dataset 1). Our model predicts a median of 14 TFs regulating each target gene and a median of 147 target genes per TF (Fig. 1c,d). 15 TFs were predicted to regulate >1,000 target genes (SI Figure 3). Importantly, TF-target gene interactions retained in our striatum-specific TRN model were enriched for genomic footprints in the adult (p = 1.4e-82) and fetal (p = 2.1e-88) brain, supporting the idea that these TF-target gene interactions reflect TF binding sites in the brain.

We defined as “TF-target gene modules” the sets of genes predicted to be direct targets of each of the 718 TFs. 135 of these 718 TF-target gene modules were enriched for a functional category from Gene Ontology (Ashburner *et al*, 2000) (FDR < 5%, adjusting for 4,624 GO terms). 337 of the 718 TF modules were enriched (p < 0.01) for genes expressed specifically in a major neuronal or non-neuronal striatal cell type (Doyle *et al*, 2008; Zhang *et al*, 2014; Dougherty *et al*, 2010), including known cell type-specific activities for both neuronal (e.g., *Npas1-3*) and glia-specific TFs (e.g., *Olig1, Olig2*) (SI Fig. 4). These results suggest that many TRN modules reflect the activities of TFs on biological processes within specific cell types.

### Prediction of core TFs associated with transcriptional changes in HD mouse models

We next sought to identify TFs that are core regulators of transcriptional changes in HD. Of the 208 mice in the RNA-seq dataset used for network reconstruction, 144 were heterozygous for a human *HTT* allele knocked into the endogenous *Htt* locus (Wheeler *et al*, 1999), and the remaining 64 mice were C57BL/6J littermate controls. Six distinct *HTT* alleles differing in the length of the poly-Q repeat were knocked in. In humans, the shortest of these alleles ‐‐ *Htt^Q20^* ‐‐ is non-pathogenic, and the remaining alleles ‐‐ *Htt^Q80^, Htt^Q92^*, *Htt^Q111^, Htt^Q140^*, and *Htt^Q175^* ‐‐ are associated with progressively earlier onset of symptoms. We used RNA-seq data from four male and four female mice of each genotype at each of three time points: 2-months-old, 6-months-old, and 10-months-old. These mouse models undergo subtle age‐ and allele-dependent changes in behavior, and all of the ages profiled precede detectable neuronal cell death (Alexandrov *et al*, 2016; Rothe *et al*, 2015; Carty *et al*, 2015).

We evaluated gene expression differences between *Htt^Q20/+^* mice and mice with each of the five pathogenic *HTT* alleles at each time point, a total of 15 comparisons. The extent of gene expression changes increased in an age‐ and Q-length-dependent fashion, with extensive overlap between the DEGs identified in each condition (Fig. 2). 8,985 genes showed some evidence of differential expression (DEGs; p < 0.01) in at least one of the 15 conditions, of which 5,132 were significant at a stringent False Discovery Rate < 1%. These results suggest that robust and replicable gene expression changes occur in the striatum of these HD mouse models at ages well before the onset of neuronal cell death or other overt pathology.

**Figure 2.**
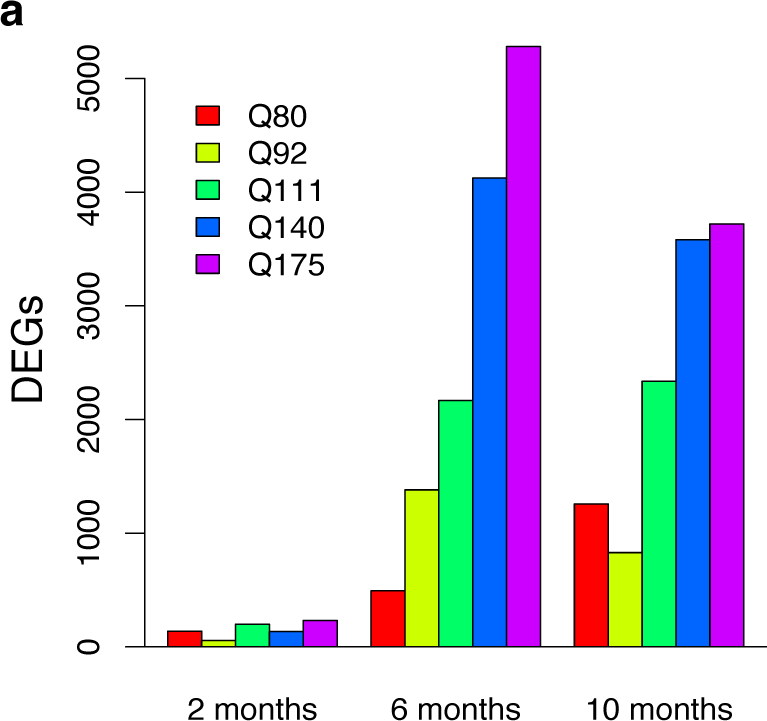
Robust changes in striatal gene expression in two-, six-, and ten-month-old HD knock-in mice. Counts of differentially expressed genes (p < 0.01) in each mouse model at each time point.

**Figure 3.**
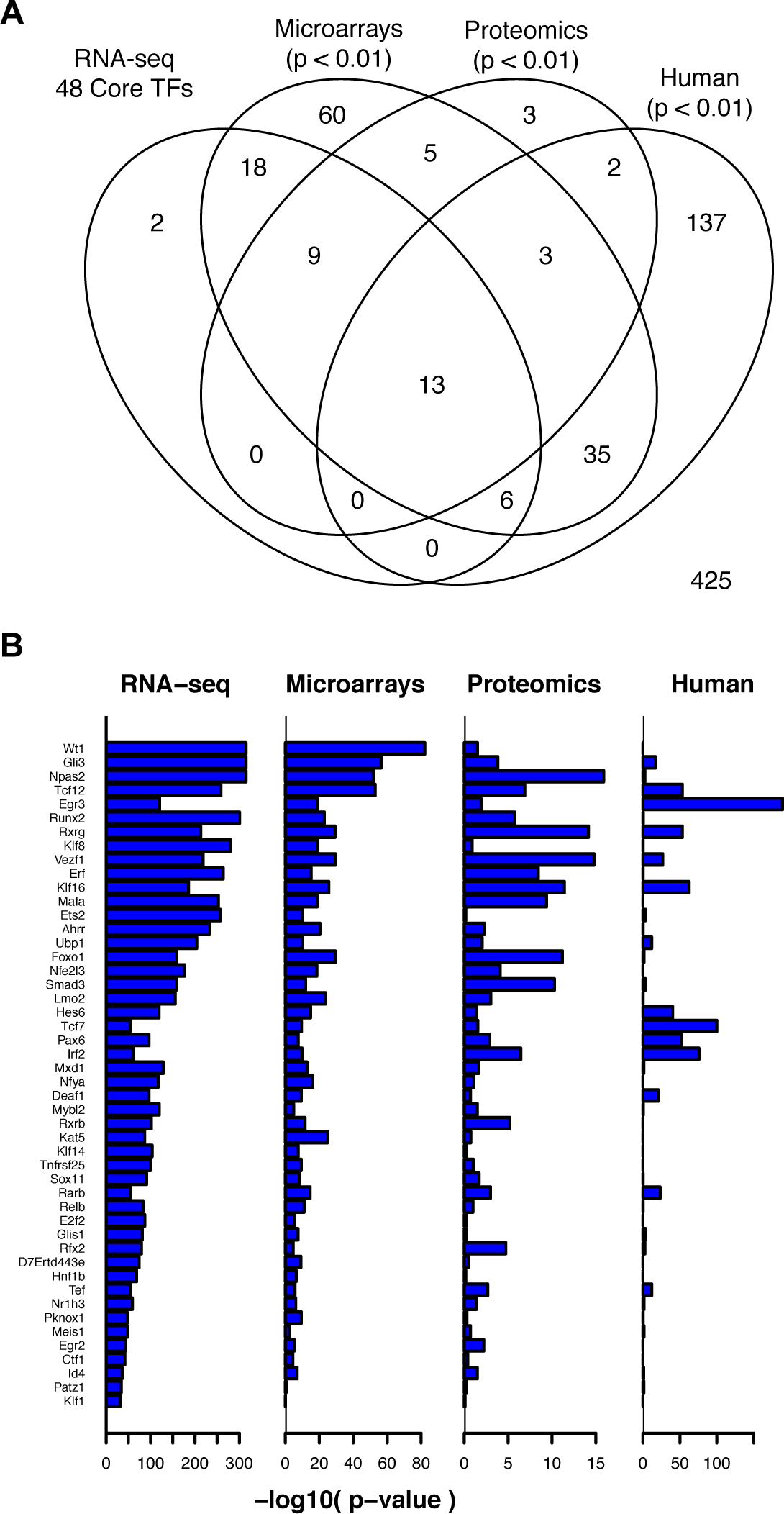
Replication of core TFs in independent datasets. a. Venn Diagram showing overlap between core regulator TF-target gene modules identified in the primary RNA-seq dataset, compared to TF-target gene modules enriched for differentially expressed genes in three independent datasets. b. −log_10_(p-values) for the strength of enrichment of each of the core regulator TF-target gene modules for differentially expressed genes in each of the four datasets.

The predicted target genes of 209 TFs were overrepresented for DEGs in at least one of the 15 conditions (3 ages x 5 mouse models; Fisher’s exact test, p < 1e-6; SI Dataset 2). Repeating this analysis in 1,000 permuted data sets indicated that enrichments at this level of significance never occurred in more than four conditions. We therefore focused on a core set of 48 TFs whose predicted target genes were overrepresented for DEGs in five or more conditions. Notably, 44 of these 48 TFs were differentially expressed (FDR < 0.01) in at least one of the 15 conditions (SI Fig. 4). We refer to these 48 TFs as core TFs.

### Replication of core TFs in independent datasets

We sought to replicate these associations by testing for enrichment of TF-target gene modules for differentially expressed genes in independent HD-related datasets. First, we conducted a meta-analysis of differentially expressed TF-target gene modules in four independent microarray gene expression profiling studies of striatal tissue from HD mouse models (Becanovic *et al*, 2010; Giles *et al*, 2012; Fossale *et al*, 2011; Kuhn *et al*, 2007). Targets of 46 of the 48 core TFs were enriched for DEGs (meta-analysis p-value < 0.01) in the microarray data. The overlap between TFs whose target genes were differentially expressed in HD vs. control mice in microarray datasets and the core TFs from our primary dataset was significantly greater than expected by chance (Fisher’s exact test: p = 5.7e-32). These results suggest that transcriptional changes in most of the core TF-target gene modules were preserved across multiple datasets and mouse models of HD.

Next, we asked whether the target genes of core TFs were also differentially abundant at the protein level. We studied quantitative proteomics data from the striatum of 64 6-month-old HD knock-in mice (Langfelder *et al*, 2016). These were a subset of the mice profiled with RNA-seq in our primary dataset. Targets of 22 of the 48 core TFs were enriched for differentially abundant proteins (Fisher’s exact test, p < 0.01). The overlap between TFs whose target genes were differentially abundant between HD vs. wildtype mice and the core regulator TFs was significantly greater than expected by chance (Fisher’s exact test: p = 5.7e-20).

Third, we asked whether target genes of these same TFs are differentially expressed in late-stage human disease. We reconstructed a TRN model for the human striatum integrating a map of TFBSs (Plaisier *et al*, 2016) based on digital genomic footprinting of 41 human cell types(Neph *et al*, 2012) with microarray gene expression profiles of post-mortem striatal tissue from 36 HD cases and 30 controls (Hodges *et al*, 2006). As in our TRN model for the mouse striatum, we fit a LASSO regression model to predict the expression of each gene in human striatum from the expression levels of TFs with predicted TFBSs within 5kb of its transcription start sites (SI Fig. 6). We studied the enrichments of TF-target gene modules from this human striatum TRN model for differentially expressed genes

We compared HD-related TF-target gene modules identified in mouse and human striatum, focusing on 616 TFs with one-to-one orthology and ≥10 predicted target genes in both the mouse and human striatum TRN models. We conducted a meta-analysis of two independent datasets from the dorsal striatum of HD cases vs. controls (Hodges *et al*, 2006; Durrenberger *et al*, 2015) to identify TF-target gene modules enriched for DEGs. Targets of 13 of the 48 core TFs from mouse striatum were over-represented among differentially expressed genes in HD cases vs. controls. This overlap was not statistically greater than expected by chance (odds ratio = 1.79; p = 0.05). However, when we considered the broader set of 209 TF-target gene modules that were enriched for differentially expressed genes in any of the 15 conditions from the primary RNA-seq dataset, we found significant overlap for TF-target gene modules that were down-regulated both in HD and in HD mouse models (28 shared TF-target gene modules; odds ratio = 3.6, p = 5.0e-5; SI Fig. 6d) and for TF-target gene modules that were up-regulated both in HD and in HD mouse models (26 shared TF-target gene modules; odds ratio = 1.8, p = 0.02; SI Fig. 6e). The striatum is heavily degraded in late-stage HD, with many dead neurons and extensive astrogliosis. Nonetheless, these results suggest that some transcriptional programs are shared between the earliest stages of molecular progression (assayed in mouse models) and late stages of human disease.

Notably, targets of 13 of the 48 core regulator TFs were enriched for differentially expressed genes in all four datasets: *Gli3*, *Irf2*, *Klf16*, *Npas2*, *Pax6*, *Rarb*, *Rfx2*, *Rxrg*, *Smad3*, *Tcf12*, *Tef*, *Ubp1*, and *Vezf1*. These 13 TFs may be especially interesting for follow-up studies.

### Biological associations of core TFs

We evaluated relationships among the 48 core TFs based on clustering and network topology. Plotting TF-to-TF regulatory interactions among the 48 core TFs (Fig. 4) revealed two distinct TF-to-TF sub-networks, characterized by numerous positive interactions within sub-networks and by fewer, mostly inhibitory interactions between sub-networks. The target genes of TFs in the first sub-network were predominantly down-regulated in HD, while the target genes of TFs in the second module were predominantly up-regulated. Hierarchical clustering of the 48 core TFs based on the expression patterns of their predicted target genes revealed similar groupings of TFs whose target genes were predominantly down‐ vs. up-regulated (Fig. 5).

**Figure 4.**
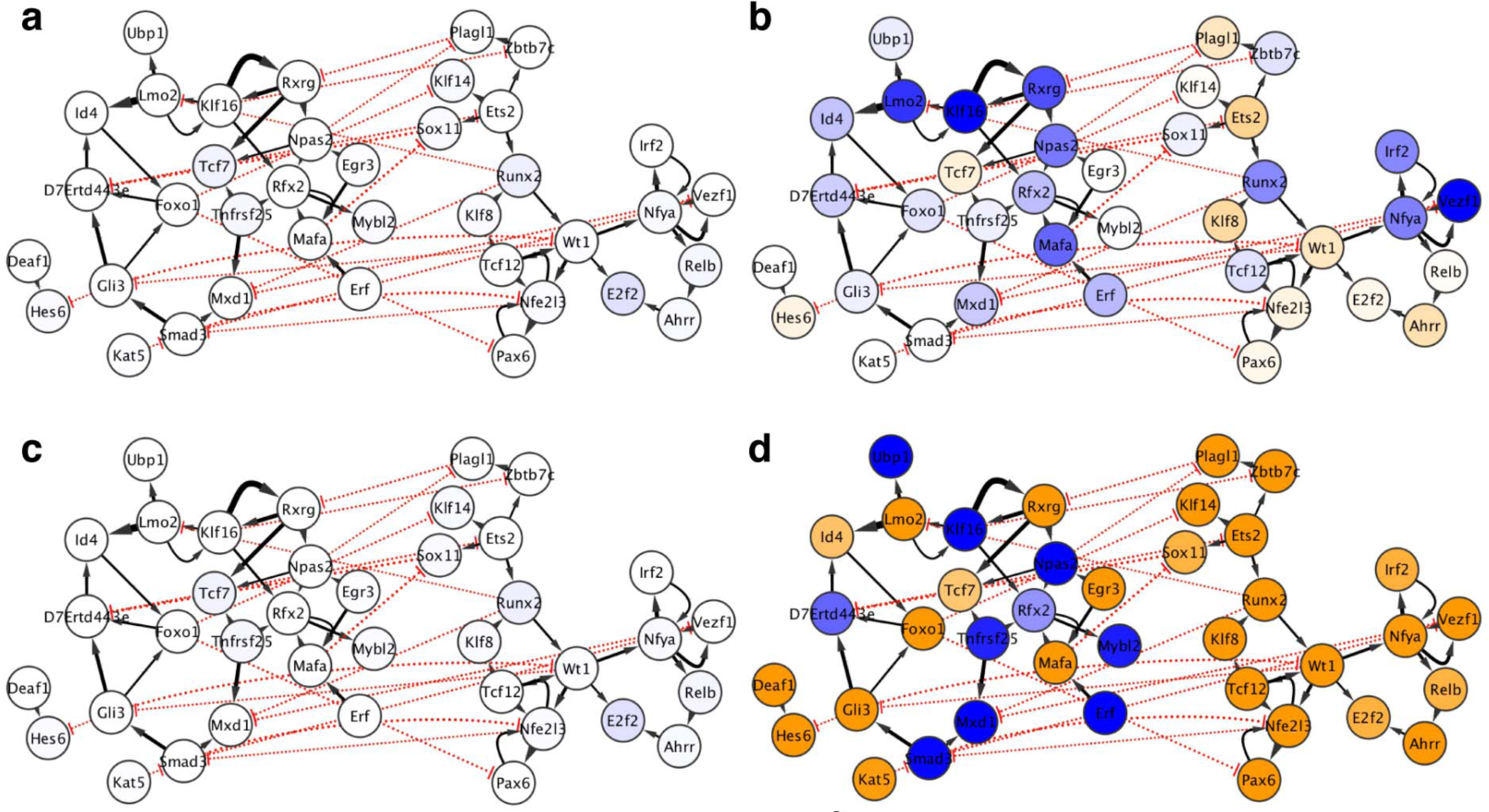
Predicted TF-to-TF interactions among 48 putative core regulators of transcriptional changes in mouse models of Huntington’s disease. Nodes and edges indicate direct regulatory interactions between TFs predicted by the mouse striatum TRN model. Solid black arrows and dotted red arrows indicate positive vs. inhibitory regulation, respectively, and the width of the line is proportional to the predicted effect size. Blue and orange shading of nodes indicates that the TF’s target genes are overrepresented for down-regulated vs. up-regulated genes in HD mouse models. If a TF’s target genes are enriched in both directions, the stronger enrichment is shown. Each panel indicates the network state in a specific condition. **a.** two-month-old *Htt^Q92/+^* mice. b. six-month-old *Htt^Q92/+^* mice. c. two-month-old *Htt^Q175/+^* mice. d. six-month-old *Htt^Q175/+^* mice.

**Figure 5.**
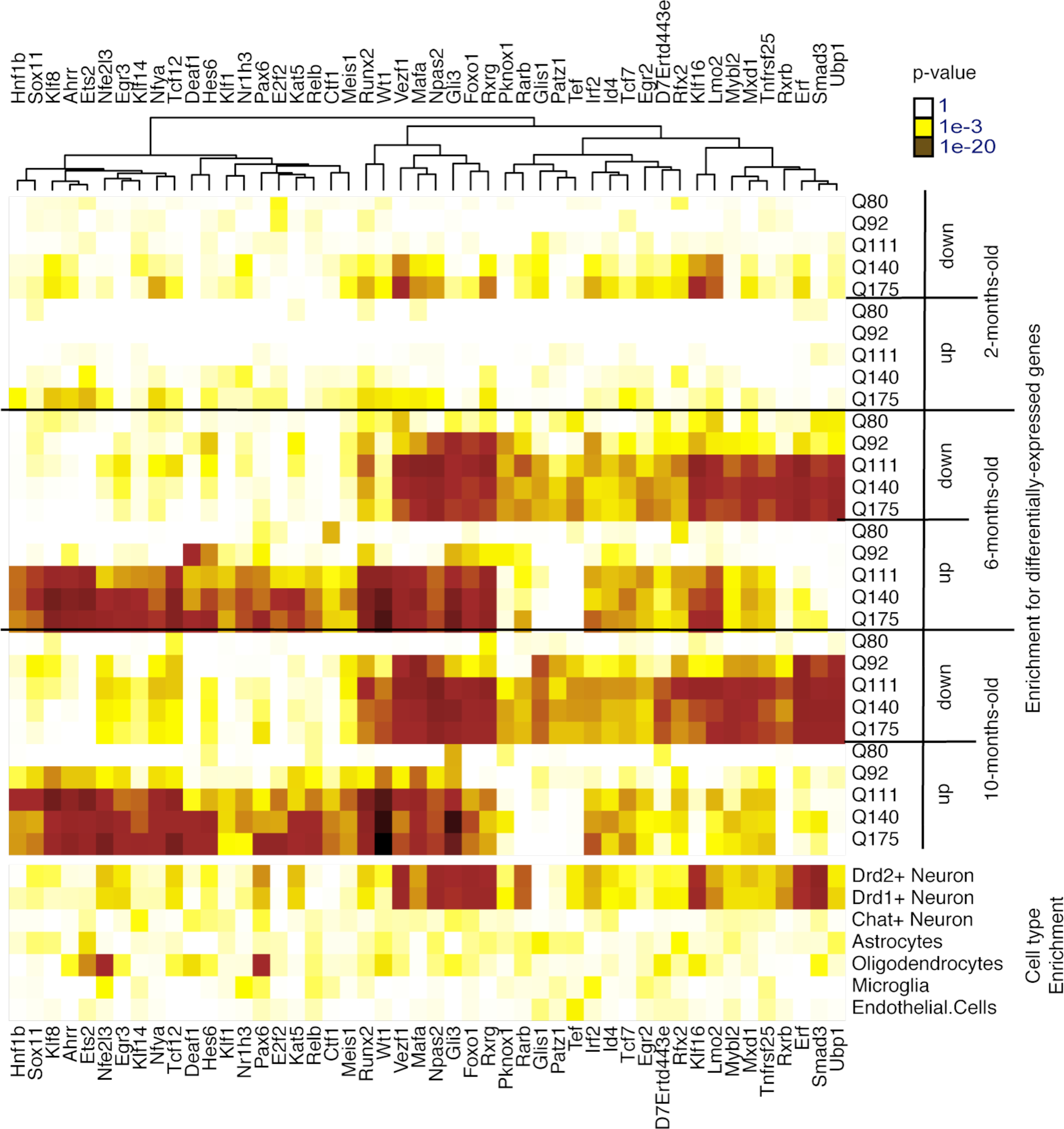
Enrichments of the 48 core TFs for differentially expressed genes in each condition and for cell type-specific genes. a. Enrichments of each TF’s target genes for down‐ and up-regulated genes for each HTT allele at each time point. b. Enrichments of each TF’s target genes for genes expressed specifically in one of seven major cell types in the mouse striatum.

We studied the predicted target genes of each core TF to characterize possible roles for these TFs in HD. Down-regulated TF-target gene modules were overrepresented for genes specifically expressed in Drd1+ and Drd2+ medium spiny neurons (Fig. 5). Functional enrichments within these modules were mostly related to synaptic function, including metal ion transmembrane transporters (targets of *Npas2*, p = 2.3e-4), voltage-gated ion channels (targets of *Mafa*, p = 8.1e-4), and protein localization to cell surface (targets of *Rxrg*, p = 1.7e-4). These network changes may be linked to synapse loss in medium spiny neurons, which is known to occur in knock-in mouse models of HD (Deng *et al*, 2013).

Some up-regulated TF-target gene modules were overrepresented for genes specifically expressed in oligodendrocytes or astrocytes, while others were overrepresented for genes specifically expressed in neurons (Fig. 5). Functional enrichments within these modules included Gene Ontology terms related to apoptosis (“positive regulation of extrinsic apoptotic signaling pathway via death domain receptors”, targets of *Wt1*, p = 1.8e-4) and DNA repair (targets of *Runx2*, “single-strand selective uracil DNA N-glycosylase activity”, p = 2.0e-4). Therefore, core TFs whose target genes were predominantly up-regulated may contribute to a variety of pathological processes both in neurons and in glia. The number of oligodendrocytes is basally increased in HD mutation carriers, while activated gliosis is thought to begin later in disease progression (Vonsattel *et al*, 1985).

An open question in the field is whether the same sequence of pathogenic events underlies disease progression in juvenile-onset HD due to *HTT* alleles with very long poly-Q tracts vs. adult-onset HD due to *HTT* alleles with relatively short poly-Q tracts. This question is of practical relevance for modeling HD in mice, since mouse models with very long *HTT* alleles are often used in research due to their faster rates of phenotypic progression within a two-year lifespan. To address this question, we evaluated overlap between TF-target gene modules activated at the earliest time points in mice with each of the five pathogenic *Htt* alleles in our dataset. In the mice with the longest *HTT* alleles ‐‐ *Htt^Q175^* and *Htt^Q140^* ‐‐ the target genes of core TFs first became enriched for differentially expressed genes in two-month-old mice. In mice with relatively short *HTT* alleles – *Htt^Q111^*, *Htt^Q92^* and *Htt^Q80^* ‐‐ target genes of core TFs became enriched for differentially expressed genes beginning in six-month-old mice. We found that eight modules – the predicted target genes of IRF2, MAFA, KLF16, LMO2, NPAS2, RUNX2, RXRG, and VEZF1 – were significantly enriched for DEGs in at least three of these five conditions (two-month-old *Htt*^Q175/+^, two-month-old *Htt^Q140/+^*, six-month-old *Htt^Q111/^*, six-month-old *Htt^Q92/+^*, and six-month-old *Htt^Q80/+^*). A limitation of this analysis is that all of the alleles used in this study are associated with juvenile onset disease, and the extent to which these results extend to adult-onset alleles remains to be detemined. Nonetheless, these results suggest that many aspects of the trajectory of transcriptional changes are shared across the *HTT* Q-lengths that have been studied. Notably, all of the TFs whose target genes were enriched for differentially expressed genes at the very earliest timepoints were enriched primarily for genes that were down-regulated in HD. Strong enrichments of TF-target gene modules for up-regulated genes occurred only at slightly later time points.

### Genome-wide characterization of SMAD3 binding sites in the mouse striatum supports a role in early gene dysregulation in HD

SMAD3 was one of 13 core TFs whose predicted target genes were overrepresented among differentially expressed genes across all four independent datasets. Progressive down-regulation of *Smad3* mRNA (Fig. 6a) and of predicted SMAD3 target genes (Fig. 5) occurred in an age‐ and *Htt*-allele-dependent fashion, beginning at or before six postnatal months.

**Figure 6.**
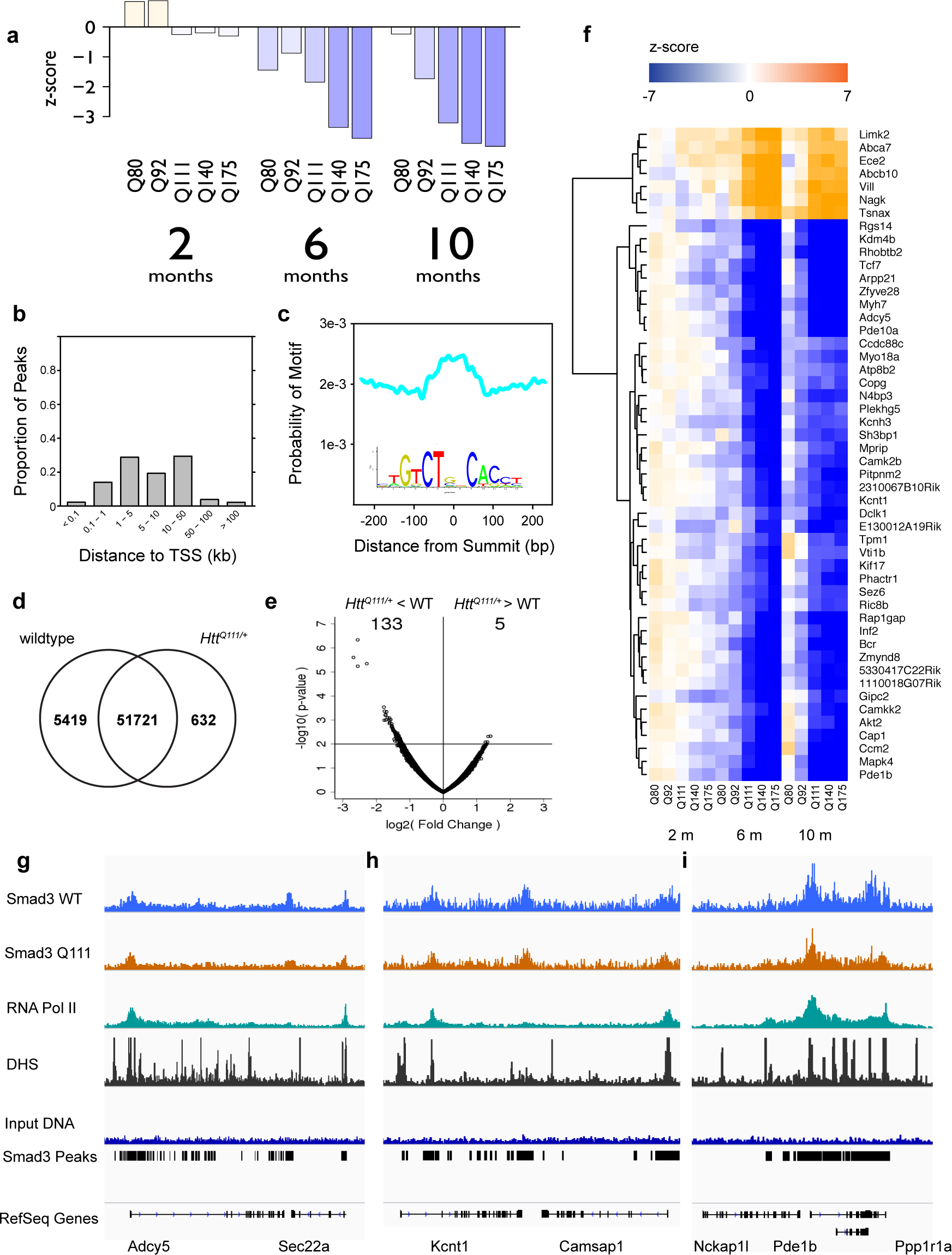
SMAD3 expression, genomic occupancy, and target gene expression in the striatum of HD mouse models. **a.** Progressive age‐ and *Htt-*allele-dependent changes in the expression of SMAD3 in mouse striatum. Bars indicate z-scores for the expression level in heterozygous mice with each pathogenic *Htt* allele compared to age-matched *Htt^Q20/+^* mice. b. Distribution of the distances of 57,772 SMAD3 peaks identified by ChIP-seq to the nearest transcription start site (TSS). c. The summits of SMAD3 peaks are enriched for the sequence motif recognized by SMAD3 (JASPAR CORE MA0513.1, shown in inset). d. Overlap between peaks identified in *Htt^Q111/+^* vs. wildtype mice. e. SMAD3 occupancy is decreased at a subset of peaks in *Htt^Q111/+^* vs. wildtype mice. x-axis and y-axis represent the log2(fold change) and – log10(p-value), respectively, for each peak region. f. Age‐ and *Htt*-allele-dependent expression patterns of the top 50 most strongly differentially expressed SMAD3 target genes. g, h, i. Genomic occupancy of SMAD3 and RNA polymerase II and accessibility of genomic DNA to DNase-I near *Adcy5* (g), *Kcnt1* (h), and *Pde1b* (i).

We characterized the binding sites of SMAD3 in the striatum of four-month-old *Htt^Q111/+^* mice and wild-type littermate controls by chromatin immunoprecipitation and deep sequencing (ChIP-seq, n=2 pooled samples per group, with each pool containing DNA from three mice). Peak-calling revealed 57,772 SMAD3 peaks (MACS2.1, FDR < 0.01 and >10 reads in at least two of the four samples; Dataset 3). 34,633 of the 57,772 SMAD3 peaks (59.9%) were located within 10kb of transcription start sites (TSSs), including at least one peak within 10kb of the TSSs for 11,727 genes (Fig. 6b). The summits of SMAD3 peaks were enriched for the SMAD2:SMAD3:SMAD4 motif (p-value = 7.2e-85; Fig. 6c). Importantly, the TSSs for 753 of the 938 computationally predicted SMAD3 target genes in our TRN model were located within 10kb of at least one ChIP-based SMAD3 binding site. This overlap was significantly greater than expected by chance (odds ratio = 4.33, p-value = 2.8e-84).

We characterized the relationship between SMAD3 occupancy and transcriptional activation by measuring the genomic occupancy of RNA Polymerase II (RNAPII) in the striatum of *Htt^Q111/+^* and wildtype mice. RNAPII occupancy is a marker of active transcription and of active transcription start sites. Occupancy of SMAD3 and of RNAPII were positively correlated, across all genomic regions (r = 0.70) and specifically within SMAD3 peaks (r = 0.71).

Similarly, we characterized the relationship between SMAD3 occupancy and chromatin accessibility, using publicly available DNase-seq of midbrain tissue from wildtype mice. 22,650 of the 57,772 SMAD3 peaks (39.2%) overlapped a DNase hypersensitive site in the midbrain. Occupancy of SMAD3 was positively correlated with DNase-I hypersensitivity across all genomic regions (r = 0.33) and specifically within SMAD3 peaks (r = 0.25).

We ranked genes from highest to lowest SMAD3 regulatory potential based on the number of SMAD3 peaks within 10kb of their transcriptional start sites. We focused on the top 837 genes with SMAD3 peak counts > 2 standard deviations above the mean. These top 837 SMAD3 target genes were enriched (FDR < 0.01) for 24 non-overlapping clusters of Gene Ontology terms (SI Table 1). These enriched GO terms prominently featured pathways related to gene regulation (“mRNA processing”, p = 4.2e-9; “histone modification”, p = 1.7e-7; “transcriptional repressor complex”, p = 3.7e-5), as well as functions more specifically related to brain function (“neuromuscular process controlling balance”, p = 1.2e-7; “brain development”, p = 1.27e-6; “neuronal cell body”, p = 2.5e-5).

We performed quantitative and qualitative analyses to compare SMAD3 occupancy in *Htt^Q111/+^* vs. wildtype mice. 51,721 of the 57,772 SMAD3 peaks (89.5%) were identified in both *Htt^Q111/+^* and wildtype mice. 5,419 peaks (9.4%) were identified only in wildtype mice, while only 632 peaks (1.1%) were identified only in *Htt^Q111/+^* mice (Fig. 6d). Quantitative analyses of differential binding with edgeR revealed four peaks whose occupancy was significantly different (FDR < 0.05) between *Htt^Q111/+^* and wildtype mice. All four of these peaks were more weakly occupied in *Htt^Q111/+^* mice. 138 peaks had nominally significant differences in occupancy between genotypes (p < 0.01). 133 of these 138 peaks (96.4%) were more weakly occupied in *Htt^Q111/+^* mice (Fig. 6e). These results suggest that SMAD3 occupancy is decreased at a subset of its binding sites in four-month-old *Htt^Q111/+^* mice.

Finally, we tested whether the top 837 SMAD3 target genes from ChIP-seq were differentially expressed in HD knock-in mice. The top 837 SMAD3 target genes from ChIP-seq were significantly overrepresented among genes that became down-regulated in the striatum of HD knock-in mice (223 down-regulated SMAD3 target genes; odds ratio = 2.0, p-value = 3.4e-15; Fig. 6f). By contrast, SMAD3 target genes were not overrepresented among genes that became up-regulated in the striatum of HD mouse models (143 up-regulated SMAD3 target genes, odds ratio = 0.92, p = 0.40). These results are consistent with our computational model, in which SMAD3 target genes were primarily down-regulated in HD knock-in mice. Therefore, SMAD3 binding is associated with down regulation in HD mouse models.

## Discussion

Here, we identified putative core TFs regulating gene expression changes in Huntington’s disease by reconstructing genome-scale transcriptional regulatory network models for the mouse and human striatum. Identifying core TFs in HD provides insights into the mechanisms of this devastating, incurable disease. This method to reconstruct models of mammalian transcriptional regulatory networks can be readily applied to find regulators underlying any trait of interest.

Our model extends prior knowledge about the TFs involved in HD. A role in HD for *Rarb* is supported by ChIP-seq and transcriptome profiling of striatal tissue from *Rarb*^-/-^ mice (Niewiadomska-Cimicka *et al*, 2016). A role in HD for *Foxo1* is supported by experimental evidence that FOXO signaling influences the vulnerability of striatal neurons to mutant Htt (Parker *et al*, 2012). A role in HD for *Relb* is supported by experimental evidence that NF-kB signaling mediates aberrant neuroinflammatory responses in HD and HD mouse models (Hsiao *et al*, 2013). Notably, microglia counts in 10-12 month *Htt^Q111/+^* mice indicate that these cells are not proliferating, suggesting that the transcriptional changes observed in our study represent a proinflammatory state, rather than microgliosis *per se*. Other predicted core TFs, including *<u>Klf16</u>* and *Rxrg*, have previously been noted among the most consistently differentially expressed genes in mouse models of HD (Seredenina & Luthi-Carter, 2012). In some cases, known functions for core TFs suggest hypotheses about their roles in HD. For instance, *Npas2* is a component of the molecular clock, so its dysfunction could contribute to circadian disturbances in HD (Morton *et al*, 2005). Notably, the predicted target genes for several TFs whose functions in HD have been studied by other investigators ‐‐ e.g., *Rest* (Zuccato *et al*, 2003), *Srebf2* (Valenza *et al*, 2005), and *Foxp1* (Tang *et al*, 2012) *–* were overrepresented for differentially expressed genes in our model, but only at later time points or more weakly than our top 48 core regulator TFs.

Our results suggest that HD involves parallel changes in distinct down‐ vs. up-regulated TF sub-networks. Targets of TFs in the down-regulated sub-network are enriched for synaptic genes and appear to be primarily neuronal. Targets of TFs in the up-regulated sub-network are enriched for stress response pathways (e.g., DNA damage repair, apoptosis). These up-regulated networks appear to involve processes occurring in both neurons and glia. Several previous studies provide independent support for synaptic changes in medium spiny neurons and of activated gliosis in HD pathogenesis (Deng *et al*, 2013; Singhrao *et al*, 1999; Hsiao *et al*, 2013).

Replication across four independent datasets revealed 13 TFs whose target genes were most consistently enriched among differentially expressed genes. We propose that these TFs should be prioritized for follow-up experiments, both to validate predicted target genes and to evaluate specific biological functions for each TF. For instance, it will be interesting to determine which (if any) of the core TFs have direct protein-protein interactions with the HTT protein and to test our model’s predictions about TF perturbations with specific aspects of HD pathology. The target genes for most of these 13 TFs were enriched for genes that were down-regulated in HD and for neuron-specific genes, consistent with the idea that pathological changes originate in medium spiny neurons.

Our ChIP-seq data confirm an association between SMAD3 binding sites and genes that are down-regulated in HD. SMAD3 is best known for its role in mediating signaling by Transforming Growth Factor-Beta (TGF-β) signaling (Kandasamy *et al*, 2011). Several recent studies have described altered TGF-β signaling in the early stages of HD (Ring *et al*, 2015; Battaglia *et al*, 2011). However, to our knowledge a role for SMAD3 has not been described. These findings suggest that intriguing possibility that agonists of TGF-β signaling could have therapeutic benefit in HD patients. Consistent with this possibility, TGF-β treatment has recently been shown to reduce apoptotic cell death in neural stem cells with expanded *HTT* polyQ tracts (Ring *et al*, 2015).

Our method to reconstruct TRNs by integrating information about TF occupancy with gene co-expression is likely to be broadly applicable, providing a strategy to optimize both mechanistic and quantitative accuracy. TRN reconstruction methods based purely on gene co-expression struggle to distinguish direct vs. indirect interactions. Physical models of TF occupancy provide poor quantitative predictions because many TF binding sites are non-functional or do not regulate the nearest gene. Our study demonstrates that integrated TRN modeling can be utilized effectively to study neurodegenerative diseases such as HD, combining data from the ENCODE project with disease specific transcriptome profiling.

## Methods

### Referenced datasets

We obtained RNA-seq and microarray gene expression profiling data from the following GEO Datasets (http://www.ncbi.nlm.nih.gov/geo/): GSE65776 (Langfelder *et al*, 2016), GSE18551 (Becanovic *et al*, 2010), GSE32417 (Giles *et al*, 2012), GSE9038 (Fossale *et al*, 2011), GSE9857 (Kuhn *et al*, 2007), GSE26927 (Durrenberger *et al*, 2015), GSE3790 (Hodges *et al*, 2006). We obtained proteomics data from the PRIDE archive (https://www.ebi.ac.uk/pride/archive/), accession PXD003442 (Langfelder *et al*, 2016). For RNA-seq data (GSE65776), we downloaded read counts and FPKM estimates, mapped to ENSEMBL gene models. For Affymetrix microarrays (GSE18551, GSE32417, GSE9038, GSE9857, GSE26927, and GSE3790) we downloaded raw image files and used the affy package in R to perform within-sample RMA normalization and between-sample quantile normalization. For proteomics data, we downloaded MaxQuant protein quantities.

### Genomic footprinting

DNase-I digestion of genomic DNA followed by deep sequencing (DNase-seq) enables the identification of genomic footprints across the complete genome. We predicted genome-wide transcription factor binding sites (TFBSs) in the mouse and human genomes based on instances of TF sequence motifs in digital genomic footprints from the ENCODE project. Short regions of genomic DNA occupied by DNA-binding proteins produce characterizes characteristic “footprints” with altered sensitivity to the DNase-I enzyme. DNase-I digestion of genomic DNA followed by deep sequencing (DNase-seq) enables the identification of genomic footprints across the complete genome.

For the human TFBS model, we used a previously described database (Plaisier *et al*, 2016) of footprints from DNase-seq of 41 cell types (Neph *et al*, 2012). For the mouse TFBS model, we downloaded digital genomic footprinting data (deep DNase-seq) for 23 mouse tissues and cell types (Yue *et al*, 2014) from the UCSC ENCODE portal on October 29, 2013: <u>ftp://hgdownload.cse.ucsc.edu/goldenPath/mm9/database/</u>. We detected footprints in each sample with Wellington (Piper *et al*, 2013), using a significance threshold, p < 1e-10. Using FIMO (Grant *et al*, 2011), we scanned the mouse genome (mm9) for instances of 2,547 motifs from TRANSFAC (Matys *et al*, 2006), JASPAR (Mathelier *et al*, 2014), UniPROBE (Hume *et al*, 2015), and high-throughput SELEX (Jolma *et al*, 2013). We intersected footprints from all tissues with motif instances to generate a genome-wide map of predicted TFBSs. A motif can be recognized by multiple TFs with similar DNA-binding domains. We assigned motifs to TF families using annotations from the TFClass database (Wingender *et al*, 2013). In total, our model included motifs recognized by 871 TFs.

### Regression-based transcriptional regulatory network models

We fit a regression model to predict the expression of each gene in mouse or human striatum, based on the expression patterns of TFs that had predicted binding sites within 5kb of that gene’s transcription start sites. We applied LASSO regularization to penalize regression coefficients and remove TFs with weak effects, using the glmnet package in R. These methods were optimized across several large transcriptomics datasets, prior to their application to the Huntsington’s disease data. To reconstruct the TRN model for mouse striatum, we used RNA-seq data from the striatum of 208 mice (Langfelder *et al*, 2016). Prior to network reconstruction, we evaluated within and between group variance and detected outlier samples using hierarchical clustering and multidimensional scaling. No major differences in variance were identified between groups, and no outlier samples were detected or removed.

We considered a variety of model parameterization during the initial model formulation. We considered elastic net regression and ridge regression as alternatives to LASSO regression. We selected LASSO based on the least falloff in performance from the training data to test sets in five-fold cross-validation. We note that when multiple TFs have correlated expression, the LASSO will generally retain only one for the final model. This feature of the LASSO has been considered advantageous, since it can eliminate indirect interactions. However, there is virtually no doubt that the TFs selected by our model underestimate the true number of TF-target gene interactions. We would only pick up dominant effects where a linear model works reasonably well. Our primary interest is ultimately in using this approach to find a relatively small number of targets based on multiple lines of evidence. We are less concerned with finding everything than in trying to make sure what we do find is as highly enriched for true positives as possible.

We also considered a variety of strategies to select an appropriate penalty parameter. For instance, we could apply an independent penalty parameter for each gene, or we could use a uniform penalty parameter across all genes. We found that optimal performance was obtained in both training data and in five-fold cross-validation when we applied a uniform penalty parameter across all genes. We assigned this penalty parameter by evaluating performance in cross-validation across a range of possible parameters for a random subset of 100 genes. For each gene, we identified the most stringent penalty such that the unfitted variance was < 1 standard error greater than the minimum unfitted variance across all the penalty parameters considered. We selected the median penalty defined by this procedure across the 100 randomly selected gene.

Not all genes’ expression can be accurately predicted based on the expression of TFs. To select genes for the final model, we evaluated the variance explained by the model in a training set consisting of 80% of the data. We selected those genes for which the model explained >50% of expression variance in the training set and carried these genes forward to a test set, consisting of the remaining 20% of genes. We found that training set performance accurately predicted test performance (r = 0.94). We therefore fit a final model for genes whose expression could be accurately predicted in the training set. The result of these procedures is a tissue-specific TRN model, predicting the TFs that regulate each gene in the striatum and assigning a positive or negative weight for each TF’s effect on that gene’s expression in the striatum.

### Enrichments of TF-target gene modules in ChIP-seq data

We downloaded ChIP-seq data from the ENCODE website (encodeproject.org, accessed August 20, 2015) for 33 mouse transcription factors included in our TRN model. We identified genes whose transcription start sites were located within 5kb of a narrowPeak in each ChIP experiment. We also downloaded a table of ChIP-to-gene annotations for 19 additional mouse TFs from the ChEA website (<u>http://amp.pharm.mssm.edu/lib/chea.jsp</u>, accessed August 6, 2015). We tested for enrichments of the target genes identified by ChIP for each of these 52 TFs to predicted TFBSs from our model.

### Enrichments of TF-target gene modules for Gene Ontology terms

We downloaded Gene Ontology (GO) annotations for mouse genes from GO.db on November 4, 2015, using the topGO R package. We extracted the genes annotated to each GO term and its children, and we used Fisher’s exact tests to characterize enrichments of TF-target gene modules for the 4,624 GO terms that contain between 10 and 500 genes.

### Enrichments of TF-target gene modules for cell type-specific genes

We characterized sets of genes expressed in each striatal cell type using gene expression profiles from purified cell types (Doyle *et al*, 2008; Zhang *et al*, 2014) and the pSI R package for Cell-type Specific Expression Analysis (Dougherty *et al*, 2010). We used Fisher’s exact tests to characterize enrichments of TF-target gene modules for genes expressed specifically in each cell type.

### Enrichments of TF-target gene modules for differentially expressed genes

We identified genes that were differentially expressed in HD vs. control samples. In the primary dataset, we compared mice with the non-pathogenic Q20 allele and mice with each of the other five alleles, separately for 2-, 6-, and 10-month-old mice. We used the edgeR R package to fit generalized linear models and test for significance of each contrast. We used Fisher’s exact tests to characterize enrichments of down-regulated genes and up-regulated genes in each condition (significance threshold for differentially expressed genes, p < 0.01) for the target genes of each TF. We considered enrichments to be statistically significant at a raw p-value threshold < 1e-6, or an adjusted p-value < 0.02 after accounting for 19,170 tests (639 TFs x 5 *Htt* alleles x 3 time points x 2 tests / condition).

To identify top TFs, accounting for non-independence among genes and conditions, we calculated an empirical false discovery rate for these enrichments. We repeated the edgeR and enrichment analyses 1,000 times with permuted sample labels. We found that no module had a p-value < 1e-6 in more than four conditions in any of the permuted datasets. Therefore, we focused on TFs whose target genes were overrepresented for differentially expressed genes in five or more conditions.

We performed similar analyses to characterize TF-target gene modules enriched for genes that were differentially expressed in replication samples. We used the limma R package to calculate differentially expressed genes in each of the four microarray studies from mouse striatum (Giles *et al*, 2012; Kuhn *et al*, 2007; Fossale *et al*, 2011; Becanovic *et al*, 2010). We calculated enrichments of the DEGs from each study for TF-target gene modules. We then combined the enrichment p-values across the four studies using Fisher’s method to produce a meta-analysis p-value for the association of each TF-target gene module in HD mouse models.

We used quantitative proteomics data from 6-month old *Htt^Q20/+^*, *Htt^Q80/+^*, *Htt^Q92/+^*, *Htt^Q111/+^*, *Htt^Q140/+^* and *Htt^Q175/+^* mice (n = 8 per group) (Langfelder *et al*, 2016). We characterized proteins whose abundance was correlated with *Htt* CAG length in the striatum of 6-month-old mice, using MaxQuant protein quantities. We then calculated enrichments of CAG-length correlated proteins (Pearson correlation, p < 0.01) for each TF-target gene module with Fisher’s exact test, separately for proteins whose abundance was positively or negatively correlated with CAG length.

We used the limma R package to fit a linear model to characterize differentially expressed genes in each of two microarray datasets (Hodges *et al*, 2006; Durrenberger *et al*, 2015) profiling dorsal striatum of HD cases vs. controls, treating sex as a covariate. We calculated enrichments of the DEGs from each study for TF-target gene modules. We then combined the enrichment p-values across the two studies using Fisher’s method to produce a meta-analysis p-value for the association of each TF-target gene module with HD.

### Mouse Breeding, Genotyping, and microdissection

The B6.*Htt^Q111/+mice^* (Strain 003456; JAX) used for the ChIP-seq study have a targeted mutation replacing a portion of mouse *Htt* (formerly *Hdh*) exon 1 with the corresponding portion of human *HTT* (formerly *IT15*) exon 1, including an expanded CAG tract (originally 109 repeats). Mice used in the present study were on the C57BL/6J inbred strain background. The targeted *Htt* allele was placed from the CD-1 background onto the C57BL/6J genetic background by selective backcrossing for more than 10 generations to the C57BL/6J strain at Jackson laboratories. Cohorts of heterozygote and wild-type littermate mice were generated by crossing B6.*Htt^Q111/+^* and B6.*Htt^+/+^* mice. Male mice were sacrificed at 122 ± 2 days of age (or 16 weeks) via a sodium phenobarbital based euthanasia solution (Fatal Plus, Henry Schein). Both hemispheres of each animal’s brain was microdissected on ice into striatum, cortex, and remaining brain regions. These tissues were snap frozen and stored in −80°C. Experiments were approved by an institutional review board in accordance with NIH animal care guidelines.

### High resolution X-ChIP-seq

We prepared duplicate ChIP samples for each antibody from four-month-old *Htt^Q111/+^* and from age-matched wildtype mice. For each ChIP preparation, chromatin DNA was prepared using the combined striatal tissue from both hemispheres of three mice. Preliminary experiments suggested that this was the minimal amount of material required to provide enough material for multiple IPs. Striata were transferred to a glass dounce on ice and homogenized in cold PBS with protease inhibitors. High-resolution X-ChIP-seq was performed as described (Skene *et al*, 2010), with slight modifications. IPs were performed using Abcam Anti-SMAD3 antibody ab28379 [ChIP grade] or Anti-RNA polymerase II CTD repeat YSPTSPS antibody [8WG16] [ChIP Grade] ab817. Sequencing libraries were prepared from the isolated ChIP DNA and from input DNA controls as previously described (Orsi *et al*, 2015). Libraries were sequenced on an Illumina HiSeq 2500 sequencer to a depth of ~17-25 million paired-end 25 bp reads per sample. Sequence reads have been deposited in GEO, accession GSE88775.

### ChIP-seq analysis

Sequencing reads were aligned to the mouse genome (mm9) using bowtie2 (Langmead & Salzberg, 2012). Peak-calling on each sample was performed with MACS v2.1 (Zhang *et al*, 2008), scaling each library to the size of the input DNA sequence library to improve comparability between samples. We retained peak regions with a significant MACS p-value (FDR < 0.01 and a read count ≥10 in at least two of the individual ChIP samples). Enrichment of the SMAD3 motif (JASPAR CORE MA0513.1) was performed with CentriMo (Bailey & Machanick, 2012), using the 250bp regions around peak summits obtained by running MACS on the combined reads from all the samples. Peaks were mapped to genes using the chipenrich R package (Welch *et al*, 2014), and genes were ranked by the number of peaks within 10kb of each gene’s transcription start sites. Gene Ontology enrichment analysis of the top SMAD3 target genes (peak counts >2 s.d. above the mean), was performed using Fisher’s exact test, using the same set of GO terms used to analyze the computationally derived TF-target gene modules. Statistical analysis of differential occupancy in *Htt^Q111/+^* vs. wildtype mice was performed with edgeR (Robinson *et al*, 2010).

### Software and Primary Data Resources

Code for analysis of gene expression, transcriptional regulatory networks, and ChIPseq data for this manuscript are publicly available in the github repository located at <u>https://github.com/seth-ament/hd-trn</u>. Bedgraph files and raw sequencing data for SMAD3 and RNA Pol2 ChIP-seq can be accessed at the GEO repository #GSE88775 prior to publication at <u>https://www.ncbi.nlm.nih.gov/geo/query/acc.cgi?token=oryzgqeerzmvdaf&acc=GSE88775</u>.

## Acknowledgements

This work was supported by a contract from the CHDI Foundation (N.D.P., Principal Investigator and J.B.C, Principal Investigator). J.R.P. is supported by a National Science Foundation Graduate Research Fellowship. J.B.C. is supported by Huntington Society of Canada New Pathways Program.

## Supplementary Information

**SI Figure 1.**
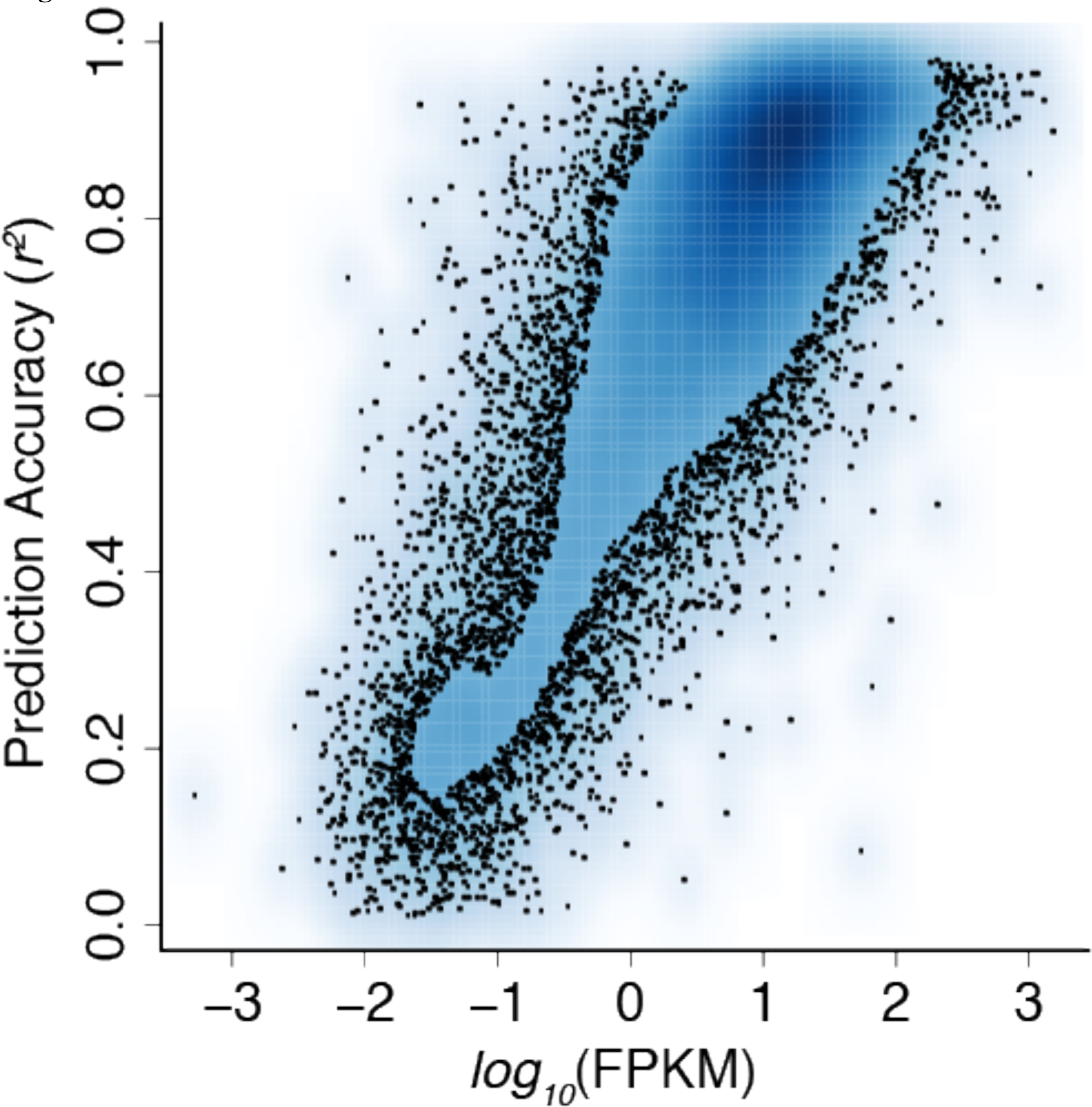
Association between TRN prediction accuracy and expression level. Each point on the scatterplot represents the mean expression level of a gene in the striatum (x-axis; fragments per kilobase million, FPKM) and the prediction accuracy for that gene in the transcriptional regulatory network model (r^2^, predicted vs. observed expression across all samples).

**SI Figure 2.**
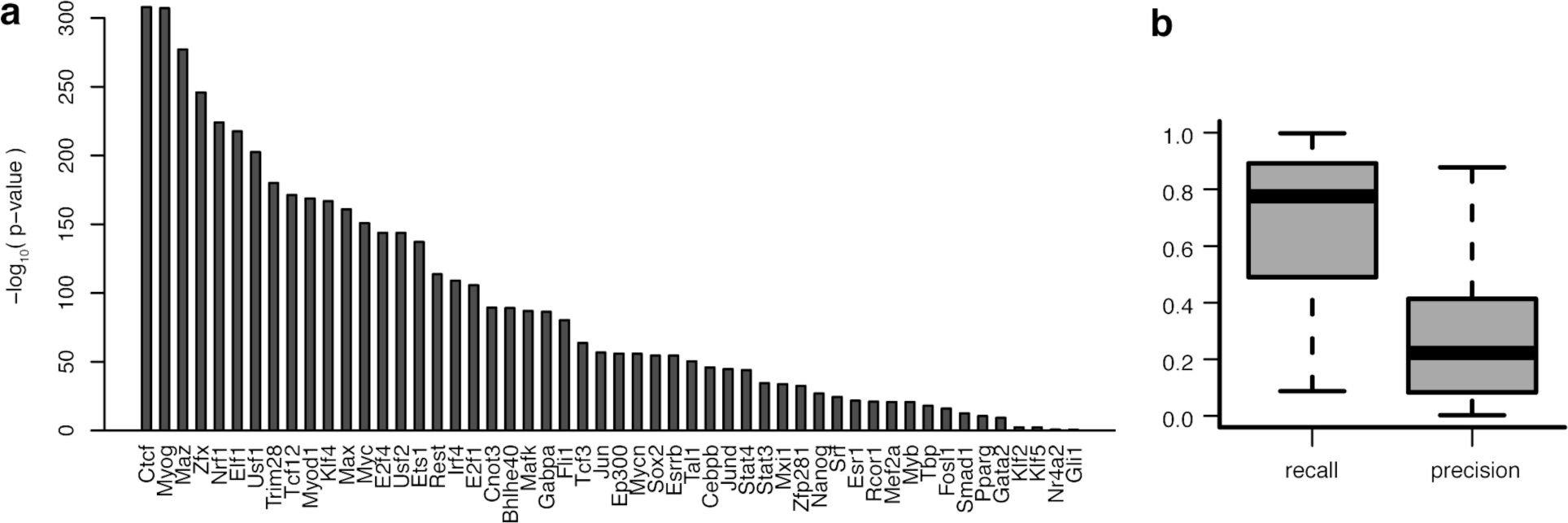
Comparison of TF binding site predictions to ChIP-seq data. For each of 52 TFs, we compared the sets of genes adjacent to predicted TF binding sites in our model to the sets of genes adjacent to observed binding sites from ChIP-seq studies. **a.** −log_10_(p-values) for overlap between modeled vs. observed gene sets (Fisher’s exact test). **b.** Distribution of recall (sensitivity) and precision (positive predictive accuracy) of the TFBS model for identifying the target genes of each TF identified by ChIP-seq.

**SI Figure 3.**
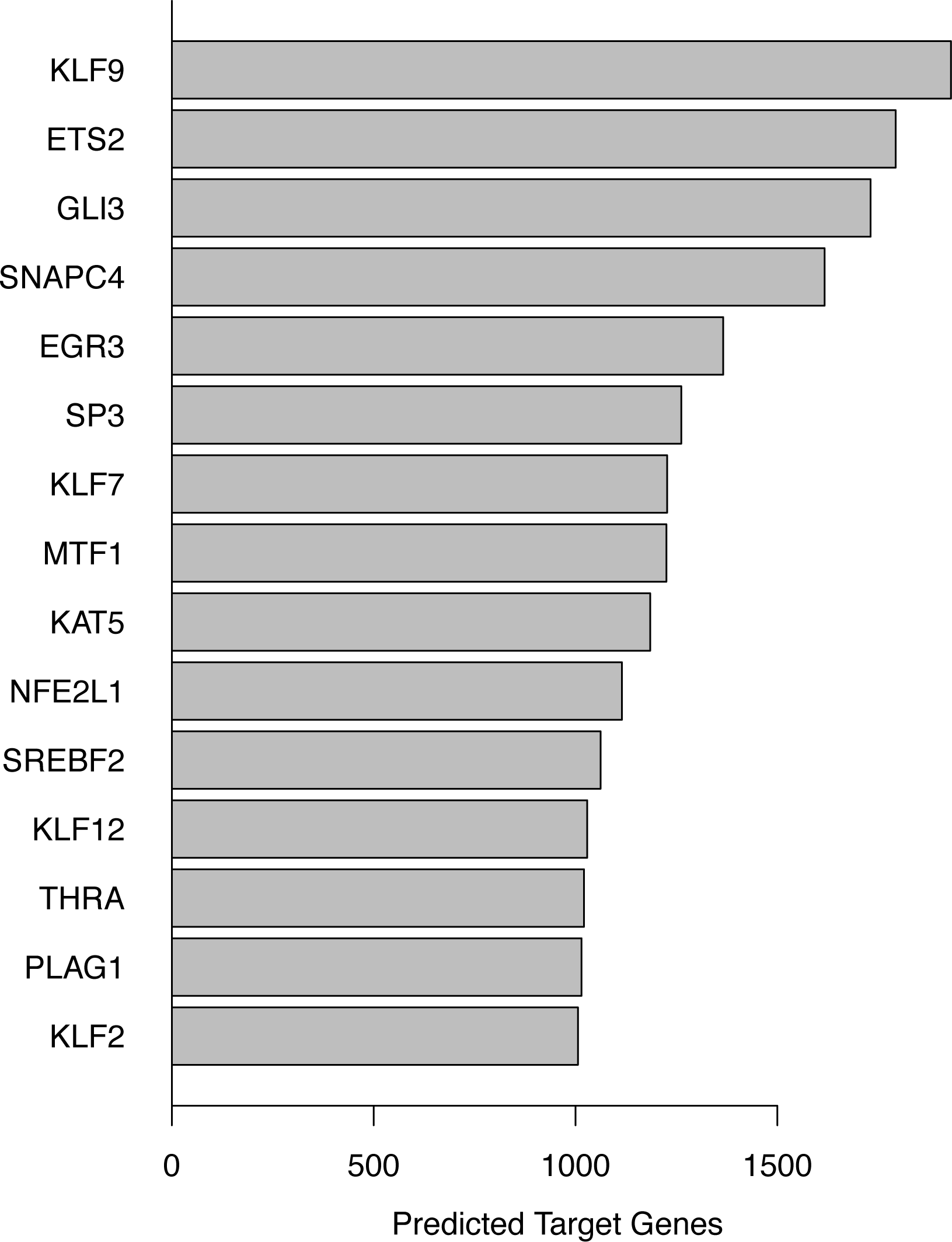
SI Figure 3. TFs with >1,000 predicted target genes. Bars indicate the number of predicted target genes for each of the 15 TFs with >1,000 predicted target genes in the TRN model for the mouse striatum.

**SI Figure 4.**
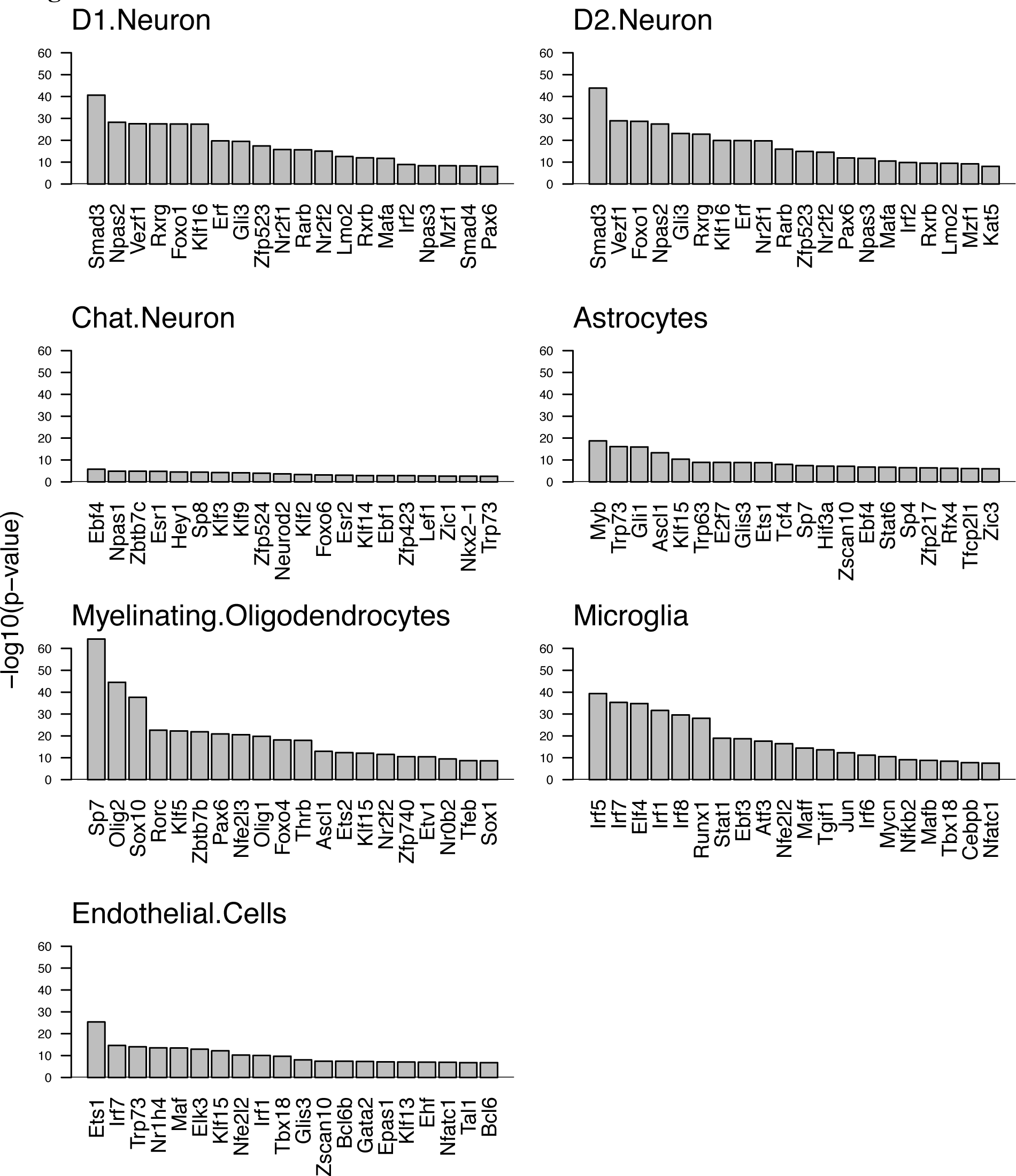
Enrichments of TF modules within each striatal cell type. Enrichments of the predicted target genes of each TF for genes expressed specifically in one of seven major cell types in the mouse striatum. The top 20 TF modules are shown for each cell type, ranked by the −log10(p-value) for the strength of enrichment in a one-sided Fisher’s exact test.

**SI Figure 5.**
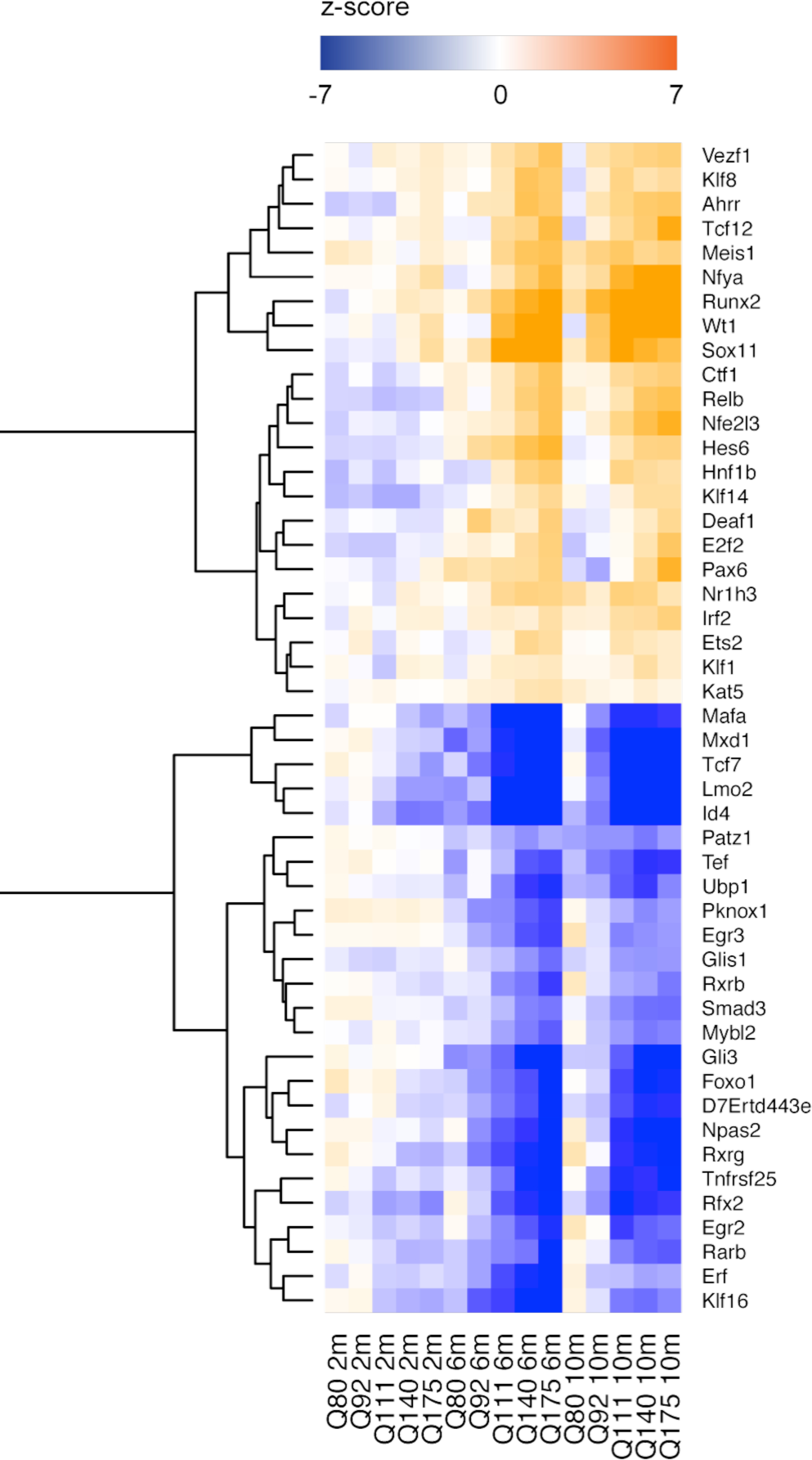
Core regulator TFs are differentially expressed in the striatum of HD CAG knock-in mice. z-scores indicate significance and direction of expression changes in each condition, relative to age-matched *Htt^Q20/+^* mice.

**SI Figure 6.**
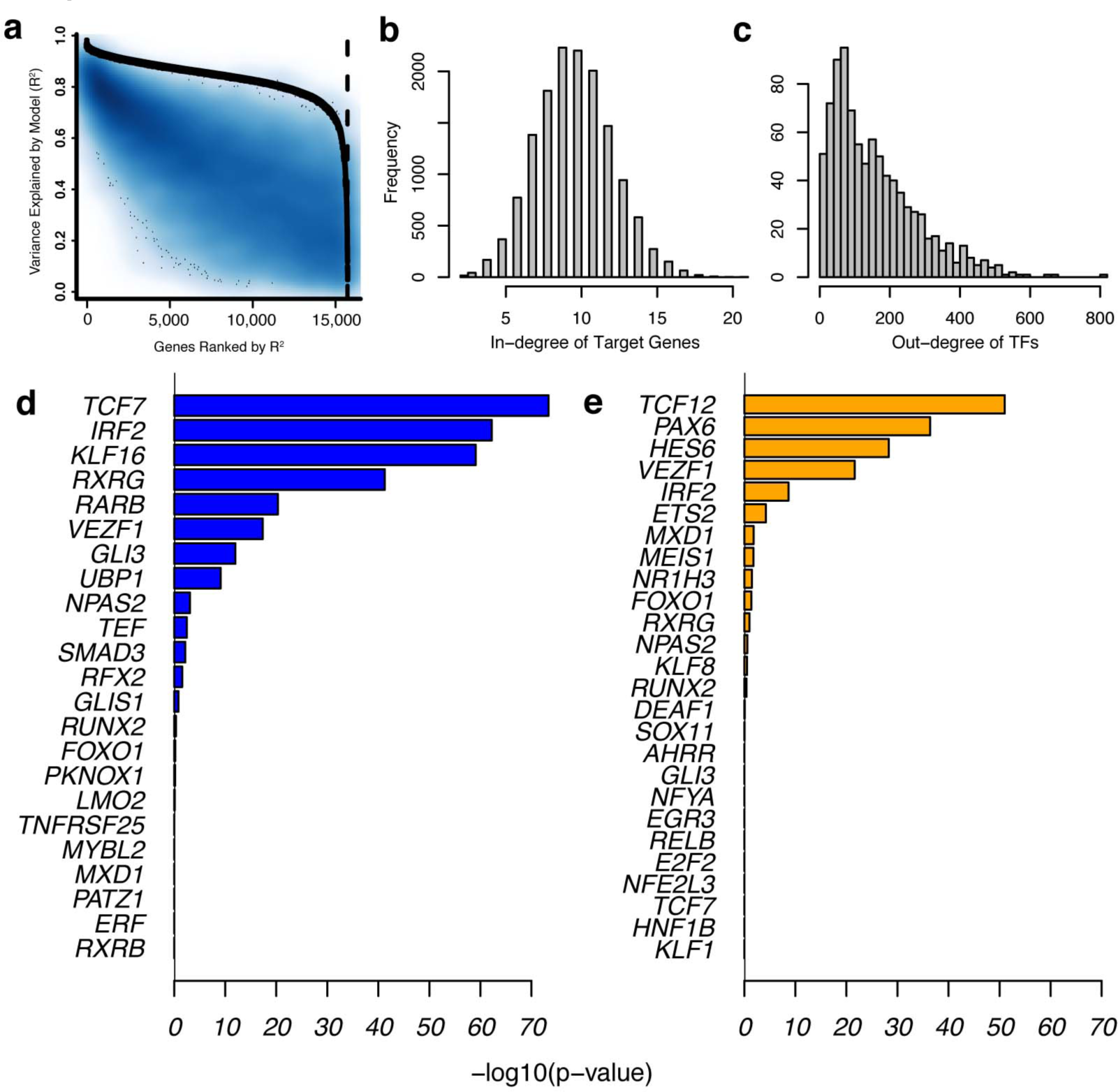
Reconstruction of a TRN model of the human striatum. a. Training (black) and test set (blue) prediction accuracy for genes in the human striatum TRN model. b. Distribution for the number of predicted regulators per target gene. c. Distribution for the number of predicted target genes per TF. d. Enrichment of down-regulated core regulator TFs identified in mouse striatum for down-regulated genes in HD cases vs. controls. e. Enrichment of up-regulated core regulator TFs identified in mouse striatum for up-regulated genes in HD cases vs. controls.

**SI Table 1.** GO enrichments of top 837 SMAD3 target genes.

| Term | Set Size | SMAD3 Targets | Odds Ratio | P-Value | FDR |
| --- | --- | --- | --- | --- | --- |
| Actin Filament-Based Process | 442 | 41 | 3.6 | 4.2E-11 | 1.9E-07 |
| mRNA Processing | 344 | 32 | 3.7 | 4.2E-09 | 9.6E-06 |
| Acting Binding | 332 | 30 | 3.6 | 2.1E-08 | 2.4E-05 |
| Neuromuscular Process Controlling Balance | 59 | 12 | 8.5 | 1.2E-07 | 9.4E-05 |
| Histone Modification | 293 | 26 | 3.6 | 1.7E-07 | 1.1E-04 |
| Brain Development | 432 | 32 | 2.8 | 1.3E-06 | 4.9E-04 |
| Chromatin Binding | 387 | 29 | 2.8 | 2.7E-06 | 9.3E-04 |
| Actin Filament-Based Movement | 65 | 11 | 6.8 | 2.8E-06 | 9.3E-04 |
| Regulation of Cell Projection Organization | 380 | 28 | 2.8 | 4.5E-06 | 1.3E-03 |
| Lamellipodium | 114 | 14 | 4.8 | 5.5E-06 | 1.4E-03 |
| Protein Serine/Threonine Kinase Activity | 409 | 29 | 2.6 | 9.9E-06 | 2.3E-03 |
| Protein Deacetylation | 49 | 9 | 7.5 | 1.1E-05 | 2.4E-03 |
| Centrosome | 350 | 25 | 2.8 | 1.4E-05 | 2.6E-03 |
| Purine Ribonucleotide Catabolic Process | 382 | 27 | 2.7 | 1.6E-05 | 3.0E-03 |
| Kinase Binding | 199 | 18 | 3.4 | 2.3E-05 | 4.0E-03 |
| Phosphoric Ester Hydrolase Activity | 374 | 26 | 2.6 | 2.5E-05 | 4.0E-03 |
| Neuronal Cell Body | 411 | 28 | 2.5 | 2.5E-05 | 4.0E-03 |
| Protein Kinase Binding | 369 | 26 | 2.6 | 2.8E-05 | 4.2E-03 |
| Cellular Protein Catabolic Process | 415 | 28 | 2.5 | 3.0E-05 | 4.2E-03 |
| Transcriptional Repressor Complex | 70 | 10 | 5.6 | 3.7E-05 | 4.4E-03 |
| Respiratory System Development | 138 | 14 | 3.8 | 5.5E-05 | 5.3E-03 |
| Kinesin Binding | 29 | 6 | 9.5 | 1.1E-04 | 9.3E-03 |
| Endocytosis | 334 | 23 | 2.6 | 1.1E-04 | 9.3E-03 |
| Negative Regulation of ERBB Signaling Pathway | 10 | 4 | 22.1 | 1.4E-04 | 9.7E-03 |

## References

Alexandrov V, Brunner D, Menalled LB, Kudwa A, Watson-Johnson J, Mazzella M, Russell I, Ruiz MC, Torello J, Sabath E, Sanchez A, Gomez M, Filipov I, Cox K, Kwan M, Ghavami A, Ramboz S, Lager B, Wheeler VC, Aaronson J, et al (2016) Large-scale phenome analysis defines a behavioral signature for Huntington’s disease genotype in mice. Nat. Biotechnol. 34: 838–44

Arlotta P, Molyneaux BJ, Jabaudon D, Yoshida Y & Macklis JD (2008) Ctip2 controls the differentiation of medium spiny neurons and the establishment of the cellular architecture of the striatum. J. Neurosci. 28: 622–32

Ashburner M, Ball CA, Blake JA, Botstein D, Butler H, Cherry JM, Davis AP, Dolinski K, Dwight SS, Eppig JT, Harris MA, Hill DP, Issel-Tarver L, Kasarskis A, Lewis S, Matese JC, Richardson JE, Ringwald M, Rubin GM & Sherlock G (2000) Gene ontology: tool for the unification of biology. The Gene Ontology Consortium. Nat. Genet. 25: 25–9

Bailey TL & Machanick P (2012) Inferring direct DNA binding from ChIP-seq. Nucleic Acids Res. 40: e128

Battaglia G, Cannella M, Riozzi B, Orobello S, Maat-Schieman ML, Aronica E, Busceti CL, Ciarmiello A, Alberti S, Amico E, Sassone J, Sipione S, Bruno V, Frati L, Nicoletti F & Squitieri F (2011) Early defect of transforming growth factor β1 formation in Huntington’s disease. J. Cell. Mol. Med. 15: 555–71

Becanovic K, Pouladi MA, Lim RS, Kuhn A, Pavlidis P, Luthi-Carter R, Hayden MR & Leavitt BR (2010) Transcriptional changes in Huntington disease identified using genome-wide expression profiling and cross-platform analysis. Hum. Mol. Genet. 19: 1438–52

Benn CL, Sun T & Sadri-Vakili G Huntingtin Modulates Transcription, Occupies Gene Promoters In Vivo, and Binds Directly to DNA in a Polyglutamine-Dependent Manner. 28:

Bonneau R, Reiss DJ, Shannon P, Facciotti M, Hood L, Baliga NS & Thorsson V (2006) The Inferelator: an algorithm for learning parsimonious regulatory networks from systems biology data sets de novo. Genome Biol. 7: R36

Carty N, Berson N, Tillack K, Thiede C, Scholz D, Kottig K, Sedaghat Y, Gabrysiak C, Yohrling G, von der Kammer H, Ebneth A, Mack V, Munoz-Sanjuan I & Kwak S (2015) Characterization of HTT inclusion size, location, and timing in the zQ175 mouse model of Huntington’s disease: an in vivo high-content imaging study. PLoS One 10: e0123527

Deng YP, Wong T, Bricker-Anthony C, Deng B & Reiner A (2013) Loss of corticostriatal and thalamostriatal synaptic terminals precedes striatal projection neuron pathology in heterozygous Q140 Huntington’s disease mice. Neurobiol. Dis. 60: 89–107

Dickey AS, Pineda V V, Tsunemi T, Liu PP, Miranda HC, Gilmore-Hall SK, Lomas N, Sampat KR, Buttgereit A, Torres M-JM, Flores AL, Arreola M, Arbez N, Akimov SS, Gaasterland T, Lazarowski ER, Ross CA, Yeo GW, Sopher BL, Magnuson GK, et al (2015) PPAR-δ is repressed in Huntington’s disease, is required for normal neuronal function and can be targeted therapeutically. Nat. Med. 22: 37–45

Dougherty JD, Schmidt EF, Nakajima M & Heintz N (2010) Analytical approaches to RNA profiling data for the identification of genes enriched in specific cells. Nucleic Acids Res. 38: 4218–30

Doyle JP, Dougherty JD, Heiman M, Schmidt EF, Stevens TR, Ma G, Bupp S, Shrestha P, Shah RD, Doughty ML, Gong S, Greengard P & Heintz N (2008) Application of a translational profiling approach for the comparative analysis of CNS cell types. Cell 135: 749–62

Durrenberger PF, Fernando FS, Kashefi SN, Bonnert TP, Seilhean D, Nait-Oumesmar B, Schmitt A, Gebicke-Haerter PJ, Falkai P, Grünblatt E, Palkovits M, Arzberger T, Kretzschmar H, Dexter DT & Reynolds R (2015) Common mechanisms in neurodegeneration and neuroinflammation: a BrainNet Europe gene expression microarray study. J. Neural Transm. 122: 1055–68

Fossale E, Seong IS, Coser KR, Shioda T, Kohane IS, Wheeler VC, Gusella JF, MacDonald ME & Lee J-M (2011) Differential effects of the Huntington’s disease CAG mutation in striatum and cerebellum are quantitative not qualitative. Hum. Mol. Genet. 20: 4258–67

Friedman J, Hastie T & Tibshirani R (2010) Regularization paths for generalized linear models via coordinate descent. J. Stat. Softw.

Friedman N, Linial M, Nachman I & Pe’er D (2000) Using Bayesian networks to analyze expression data. J. Comput. Biol. 7: 601–20

Gerstein MB, Kundaje A, Hariharan M, Landt SG, Yan K-K, Cheng C, Mu XJ, Khurana E, Rozowsky J, Alexander R, Min R, Alves P, Abyzov A, Addleman N, Bhardwaj N, Boyle AP, Cayting P, Charos A, Chen DZ, Cheng Y, et al (2012) Architecture of the human regulatory network derived from ENCODE data. Nature 489: 91–100

Giles P, Elliston L, Higgs G V, Brooks SP, Dunnett SB & Jones L (2012) Longitudinal analysis of gene expression and behaviour in the HdhQ150 mouse model of Huntington’s disease. Brain Res. Bull. 88: 199–209

Grant CE, Bailey TL & Noble WS (2011) FIMO: scanning for occurrences of a given motif. Bioinformatics 27: 1017–8

Haury A-C, Mordelet F, Vera-Licona P & Vert J-P (2012) TIGRESS: Trustful Inference of Gene REgulation using Stability Selection. BMC Syst. Biol. 6: 145

Hodges A, Strand AD, Aragaki AK, Kuhn A, Sengstag T, Hughes G, Elliston LA, Hartog C, Goldstein DR, Thu D, Hollingsworth ZR, Collin F, Synek B, Holmans PA, Young AB, Wexler NS, Delorenzi M, Kooperberg C, Augood SJ, Faull RLM, et al (2006) Regional and cellular gene expression changes in human Huntington’s disease brain. Hum. Mol. Genet. 15: 965–77

Hsiao H-Y, Chen Y-C, Chen H-M, Tu P-H & Chern Y (2013a) A critical role of astrocyte mediated nuclear factor-?B-dependent inflammation in Huntington’s disease. Hum. Mol. Genet. 22: 1826–42

Hsiao H-Y, Chen Y-C, Chen H-M, Tu P-H & Chern Y (2013b) A critical role of astrocyte mediated nuclear factor-?B-dependent inflammation in Huntington’s disease. Hum. Mol. Genet. 22: 1826–42

Hume MA, Barrera LA, Gisselbrecht SS & Bulyk ML (2015) UniPROBE, update 2015: new tools and content for the online database of protein-binding microarray data on protein DNA interactions. Nucleic Acids Res. 43: D117-22

Huntingtin interacts with REST/NRSF to modulate the transcription of NRSE-controlled neuronal genes (2003) Nat. …

Huynh-Thu VA, Irrthum A, Wehenkel L & Geurts P (2010) Inferring regulatory networks from expression data using tree-based methods. PLoS One 5:

Jolma A, Yan J, Whitington T, Toivonen J, Nitta KR, Rastas P, Morgunova E, Enge M, Taipale M, Wei G, Palin K, Vaquerizas JM, Vincentelli R, Luscombe NM, Hughes TR, Lemaire P, Ukkonen E, Kivioja T & Taipale J (2013) DNA-binding specificities of human transcription factors. Cell 152: 327–39

Kandasamy M, Reilmann R, Winkler J, Bogdahn U & Aigner L (2011) Transforming Growth Factor-Beta Signaling in the Neural Stem Cell Niche: A Therapeutic Target for Huntington’s Disease. Neurol. Res. Int. 2011: 124256

Kuhn A, Goldstein DR, Hodges A, Strand AD, Sengstag T, Kooperberg C, Becanovic K, Pouladi MA, Sathasivam K, Cha J-HJ, Hannan AJ, Hayden MR, Leavitt BR, Dunnett SB, Ferrante RJ, Albin R, Shelbourne P, Delorenzi M, Augood SJ, Faull RLM, et al (2007) Mutant huntingtin’s effects on striatal gene expression in mice recapitulate changes observed in human Huntington’s disease brain and do not differ with mutant huntingtin length or wild type huntingtin dosage. Hum. Mol. Genet. 16: 1845–61

Lachmann A, Xu H, Krishnan J, Berger SI, Mazloom AR & Ma’ayan A (2010) ChEA: transcription factor regulation inferred from integrating genome-wide ChIP-X experiments. Bioinformatics 26: 2438–44

Langfelder P, Cantle JP, Chatzopoulou D, Wang N, Gao F, Al-Ramahi I, Lu X-H, Ramos EM, El-Zein K, Zhao Y, Deverasetty S, Tebbe A, Schaab C, Lavery DJ, Howland D, Kwak S, Botas J, Aaronson JS, Rosinski J, Coppola G, et al (2016) Integrated genomics and proteomics define huntingtin CAG length-dependent networks in mice. Nat. Neurosci. 19: 623–33

Langmead B & Salzberg SL (2012) Fast gapped-read alignment with Bowtie 2. Nat. Methods 9: 357–9

Li L, Liu H, Dong P, Li D, Legant WR, Grimm JB, Lavis LD, Betzig E, Tjian R & Liu Z (2016) Real-time imaging of Huntingtin aggregates diverting target search and gene transcription. Elife 5: 1–29

Luthi-Carter R, Strand A, Peters NL, Solano SM, Hollingsworth ZR, Menon AS, Frey AS, Spektor BS, Penney EB, Schilling G, Ross CA, Borchelt DR, Tapscott SJ, Young AB, Cha JH & Olson JM (2000) Decreased expression of striatal signaling genes in a mouse model of Huntington’s disease. Hum. Mol. Genet. 9: 1259–71

MacDonald M, Ambrose C & Duyao M (1993) A novel gene containing a trinucleotide repeat that is expanded and unstable on Huntington’s disease chromosomes. Cell

Marbach D, Costello JC, Küffner R, Vega NM, Prill RJ, Camacho DM, Allison KR, Kellis M, Collins JJ & Stolovitzky G (2012) Wisdom of crowds for robust gene network inference. Nat. Methods 9: 796–804

Margolin AA, Nemenman I, Basso K, Wiggins C, Stolovitzky G, Dalla Favera R & Califano A (2006) ARACNE: an algorithm for the reconstruction of gene regulatory networks in a mammalian cellular context. BMC Bioinformatics 7 Suppl 1: S7

Mathelier A, Zhao X, Zhang AW, Parcy F, Worsley-Hunt R, Arenillas DJ, Buchman S, Chen C, Chou A, Ienasescu H, Lim J, Shyr C, Tan G, Zhou M, Lenhard B, Sandelin A & Wasserman WW (2014) JASPAR 2014: an extensively expanded and updated open-access database of transcription factor binding profiles. Nucleic Acids Res. 42: D142-7

Matys V, Kel-Margoulis O V, Fricke E, Liebich I, Land S, Barre-Dirrie A, Reuter I, Chekmenev D, Krull M, Hornischer K, Voss N, Stegmaier P, Lewicki-Potapov B, Saxel H, Kel AE & Wingender E (2006) TRANSFAC and its module TRANSCompel: transcriptional gene regulation in eukaryotes. Nucleic Acids Res. 34: D108-10

Morton AJ, Wood NI, Hastings MH, Hurelbrink C, Barker RA & Maywood ES (2005) Disintegration of the sleep-wake cycle and circadian timing in Huntington’s disease. J. Neurosci. 25: 157–63

Neph S, Vierstra J, Stergachis AB, Reynolds AP, Haugen E, Vernot B, Thurman RE, John S, Sandstrom R, Johnson AK, Maurano MT, Humbert R, Rynes E, Wang H, Vong S, Lee K, Bates D, Diegel M, Roach V, Dunn D, et al (2012) An expansive human regulatory lexicon encoded in transcription factor footprints. Nature 489: 83–90

Niewiadomska-Cimicka A, Krzyżosiak A, Ye T, Podleśny-Drabiniok A, Dembélé D, Dollé P & Krężel W (2016) Genome-wide Analysis of RARβ Transcriptional Targets in Mouse Striatum Links Retinoic Acid Signaling with Huntington’s Disease and Other Neurodegenerative Disorders. Mol. Neurobiol.

Orsi GA, Kasinathan S, Zentner GE, Henikoff S & Ahmad K (2015) Mapping regulatory factors by immunoprecipitation from native chromatin. Curr. Protoc. Mol. Biol. 110: 21.31.1-25

Parker JA, Vazquez-Manrique RP, Tourette C, Farina F, Offner N, Mukhopadhyay A, Orfila A-M, Darbois A, Menet S, Tissenbaum HA & Neri C (2012) Integration of β-catenin, sirtuin, and FOXO signaling protects from mutant huntingtin toxicity. J. Neurosci. 32: 12630–40

Piper J, Elze MC, Cauchy P, Cockerill PN, Bonifer C & Ott S (2013) Wellington: a novel method for the accurate identification of digital genomic footprints from DNase-seq data. Nucleic Acids Res. 41: e201

Plaisier CL, O’Brien S, Bernard B, Reynolds S, Simon Z, Toledo CM, Ding Y, Reiss DJ, Paddison PJ & Baliga NS (2016) Causal Mechanistic Regulatory Network for Glioblastoma Deciphered Using Systems Genetics Network Analysis. Cell Syst.

Reiss DJ, Plaisier CL, Wu W-J & Baliga NS (2015) cMonkey2: Automated, systematic, integrated detection of co-regulated gene modules for any organism. Nucleic Acids Res. 43: e87

Ring KL, An MC, Zhang N, O’Brien RN, Ramos EM, Gao F, Atwood R, Bailus BJ, Melov S, Mooney SD, Coppola G & Ellerby LM (2015) Genomic Analysis Reveals Disruption of Striatal Neuronal Development and Therapeutic Targets in Human Huntington’s Disease Neural Stem Cells. Stem cell reports 5: 1023–38

Robinson MD, McCarthy DJ & Smyth GK (2010) edgeR: a Bioconductor package for differential expression analysis of digital gene expression data. Bioinformatics 26: 139–40

Rothe T, Deliano M, Wójtowicz AM, Dvorzhak A, Harnack D, Paul S, Vagner T, Melnick I, Stark H & Grantyn R (2015) Pathological gamma oscillations, impaired dopamine release, synapse loss and reduced dynamic range of unitary glutamatergic synaptic transmission in the striatum of hypokinetic Q175 Huntington mice. Neuroscience 311: 519–38

Seong IS, Woda JM, Song J-J, Lloret A, Abeyrathne PD, Woo CJ, Gregory G, Lee J-M, Wheeler VC, Walz T, Kingston RE, Gusella JF, Conlon RA & MacDonald ME (2010) Huntingtin facilitates polycomb repressive complex 2. Hum. Mol. Genet. 19: 573–83

Seredenina T & Luthi-Carter R (2012) What have we learned from gene expression profiles in Huntington’s disease? Neurobiol. Dis. 45: 83–98

Shirasaki DI, Greiner ER, Al-Ramahi I, Gray M, Boontheung P, Geschwind DH, Botas J, Coppola G, Horvath S, Loo JA & Yang XW (2012) Network organization of the huntingtin proteomic interactome in mammalian brain. Neuron 75: 41–57

Singhrao SK, Neal JW, Morgan BP & Gasque P (1999) Increased complement biosynthesis by microglia and complement activation on neurons in Huntington’s disease. Exp. Neurol. 159: 362–76

Skene PJ, Illingworth RS, Webb S, Kerr ARW, James KD, Turner DJ, Andrews R & Bird AP (2010) Neuronal MeCP2 is expressed at near histone-octamer levels and globally alters the chromatin state. Mol. Cell 37: 457–68

Tabrizi SJ, Scahill RI, Owen G, Durr A, Leavitt BR, Roos RA, Borowsky B, Landwehrmeyer B, Frost C, Johnson H, Craufurd D, Reilmann R, Stout JC, Langbehn DR & TRACK-HD Investigators (2013) Predictors of phenotypic progression and disease onset in premanifest and early-stage Huntington’s disease in the TRACK-HD study: analysis of 36-month observational data. Lancet. Neurol. 12: 637–49

Tang B, Becanovic K, Desplats PA, Spencer B, Hill AM, Connolly C, Masliah E, Leavitt BR & Thomas EA (2012) Forkhead box protein p1 is a transcriptional repressor of immune signaling in the CNS: implications for transcriptional dysregulation in Huntington disease. Hum. Mol. Genet. 21: 3097–111

Thomas EA, Coppola G & Desplats PA The HDAC inhibitor 4b ameliorates the disease phenotype and transcriptional abnormalities in Huntington’s disease transgenic mice. 105:

Tibshirani R (1996) Regression shrinkage and selection via the lasso. J. R. Stat. Soc. Ser. B ( …

Valenza M, Rigamonti D, Goffredo D, Zuccato C, Fenu S, Jamot L, Strand A, Tarditi A, Woodman B, Racchi M, Mariotti C, Di Donato S, Corsini A, Bates G, Pruss R, Olson JM, Sipione S, Tartari M & Cattaneo E (2005) Dysfunction of the cholesterol biosynthetic pathway in Huntington’s disease. J. Neurosci. 25: 9932–9

Vonsattel JP, Myers RH, Stevens TJ, Ferrante RJ, Bird ED & Richardson EP (1985) Neuropathological classification of Huntington’s disease. J. Neuropathol. Exp. Neurol. 44: 559–77

Welch RP, Lee C, Imbriano PM, Patil S, Weymouth TE, Smith RA, Scott LJ & Sartor MA (2014) ChIP-Enrich: gene set enrichment testing for ChIP-seq data. Nucleic Acids Res. 42: e105

Wheeler VC, Auerbach W, White JK, Srinidhi J, Auerbach A, Ryan A, Duyao MP, Vrbanac V, Weaver M, Gusella JF, Joyner AL & MacDonald ME (1999) Length-dependent gametic CAG repeat instability in the Huntington’s disease knock-in mouse. Hum. Mol. Genet. 8: 115–22

Wheeler VC, White JK, Gutekunst CA, Vrbanac V, Weaver M, Li XJ, Li SH, Yi H, Vonsattel JP, Gusella JF, Hersch S, Auerbach W, Joyner AL & MacDonald ME (2000) Long glutamine tracts cause nuclear localization of a novel form of huntingtin in medium spiny striatal neurons in HdhQ92 and HdhQ111 knock-in mice. Hum. Mol. Genet. 9: 503–13

Wingender E, Schoeps T & Dönitz J (2013) TFClass: an expandable hierarchical classification of human transcription factors. Nucleic Acids Res. 41: D165-70

Yue F, Cheng Y, Breschi A, Vierstra J, Wu W, Ryba T, Sandstrom R, Ma Z, Davis C, Pope BD, Shen Y, Pervouchine DD, Djebali S, Thurman RE, Kaul R, Rynes E, Kirilusha A, Marinov GK, Williams BA, Trout D, et al (2014) A comparative encyclopedia of DNA elements in the mouse genome. Nature 515: 355–64

Zhang Y, Chen K, Sloan SA, Bennett ML, Scholze AR, O’Keeffe S, Phatnani HP, Guarnieri P, Caneda C, Ruderisch N, Deng S, Liddelow SA, Zhang C, Daneman R, Maniatis T, Barres BA & Wu JQ (2014) An RNA-Sequencing Transcriptome and Splicing Database of Glia, Neurons, and Vascular Cells of the Cerebral Cortex. J. Neurosci. 34: 11929–47

Zhang Y, Liu T, Meyer CA, Eeckhoute J, Johnson DS, Bernstein BE, Nusbaum C, Myers RM, Brown M, Li W & Liu XS (2008) Model-based analysis of ChIP-Seq (MACS). Genome Biol. 9: R137

Zuccato C, Belyaev N, Conforti P, Ooi L, Tartari M, Papadimou E, MacDonald M, Fossale E, Zeitlin S, Buckley N & Cattaneo E (2007) Widespread disruption of repressor element-1 silencing transcription factor/neuron-restrictive silencer factor occupancy at its target genes in Huntington’s disease. J. Neurosci. 27: 6972–83

Zuccato C, Tartari M, Crotti A, Goffredo D, Valenza M, Conti L, Cataudella T, Leavitt BR, Hayden MR, Timmusk T, Rigamonti D & Cattaneo E (2003) Huntingtin interacts with REST/NRSF to modulate the transcription of NRSE-controlled neuronal genes. Nat. Genet. 35: 76–83

